# An allosteric interaction controls the activation mechanism of SHP2 tyrosine phosphatase

**DOI:** 10.1101/2020.03.24.004911

**Authors:** Massimiliano Anselmi, Jochen S. Hub

**Affiliations:** Institute for Microbiology and Genetics, Georg-August-Universität Göttingen, 37077 Göttingen, Germany; Theoretical Physics and Center for Biophysics, Saarland University, Campus E2.6, 66123 Saarbrücken, Germany

**Keywords:** Molecular dynamics simulations, Phosphatases, Protein dynamics, Protein function, Allosteric regulation

## Abstract

SHP2 is a protein tyrosine phosphatase (PTP) involved in multiple signaling pathways. Mutations of SHP2 can result in Noonan syndrome or pediatric malignancies. Inhibition of wild-type SHP2 represents a novel strategy against several cancers. SHP2 is activated by binding of a phosphopeptide to the N-SH2 domain of SHP2, thereby favoring dissociation of the N-SH2 domain and exposing the active site on the PTP domain. The conformational transitions controlling ligand affinity and PTP dissociation remain poorly understood. Using molecular simulations, we revealed an allosteric interaction restraining the N-SH2 domain into a SHP2-activating and a stabilizing state. Only ligands selecting for the activating N-SH2 conformation, depending on ligand sequence and binding mode, are effective activators. We validate the model of SHP2 activation by rationalizing modified basal activity and responsiveness to ligand stimulation of several N-SH2 variants. This study provides mechanistic insight into SHP2 activation and may open routes for SHP2 regulation.

## INTRODUCTION

Reversible tyrosine phosphorylation is a post-translational modification reciprocally controlled by protein-tyrosine kinases (PTKs) and phosphatases (PTPs), playing key roles in regulating cell proliferation, differentiation, migration, apoptosis as well as cell–cell communication. Abnormal control of tyrosyl phosphorylation can result in various human diseases, including cancer (1, 2).

Among PTPs, the SH2 domain-containing phosphatase SHP2 is a regulator of signaling downstream of several cell surface receptors, functioning as positive or negative modulator in multiple signaling pathways (3). Notably, SHP2 is required for full activation of the RAS/MAPK signaling cascade, and dominantly acting mutations of *PTPN11,* the gene encoding SHP2, cause developmental disorders (i.e., Noonan syndrome (4, 5, 6, 7) and Noonan syndrome with multiple lentigines (6, 8)). Somatic mutations in *PTPN11* contribute to childhood malignancies, among which juvenile myelomonocytic leukemia (JMML) represents the archetypal disorder resulting from RAS signaling upregulation (5, 9, 10, 11, 12). Recent studies suggested SHP2 inhibition as a promising strategy for treating a large class of receptor tyrosine kinase-driven cancers (13) and for combating resistance to targeted anticancer therapies (14, 15, 16). In addition, SHP2 plays a role in *Helicobacter pylori*-induced gastric cancer mediated by activation by the bacterial protein CagA (17), and SHP2 is responsible for the suppression of T-cell activation by programmed cell death-1 (PD-1), a receptor hijacked by tumor cells for evading the immune response (18). All these recent discoveries indicate SHP2 as an attractive target in future anti-cancer therapies (19, 20, 21).

The structure of SHP2 includes two tandemly-arranged Src homology 2 domains, called N-SH2 and C-SH2, followed by the catalytic PTP domain, and a C-terminal tail with a still poorly characterized function. The SH2 domains are recognition elements that allow SHP2 to bind to signaling partners containing a phosphorylated tyrosine (pY) (22). Crystal structures have revealed an allosteric regulation of SHP2 activity (23). Namely, under basal conditions, SHP2 is in an autoinhibited state where the blocking loop of the N-SH2 domain occludes the catalytic site of the PTP domain. Association of SHP2 to its binding partners via the SH2 domains favors the release of the autoinhibitory N-SH2–PTP interactions, rendering the catalytic site accessible to substrates (23, 24, 25). Whereas this mode of regulation is widely accepted, the molecular mechanisms underlying SHP2 activation, binding partner recognition, and allostery of the N-SH2 domain are not well understood.

Atomic structures are available for the autoinhibited, closed state of SHP2 (23, 26), and for the isolated N-SH2 domain, either with a bound phosphopeptide or in its apo form (24). In addition, a recent structure of the “open” state (obtained for the basally active, leukemia-associated E76K mutant) revealed an alternative relative arrangement of N-SH2 and PTP domains, with the active site within the PTP domain exposed to the solvent (25). According to these structures, N-SH2 undergoes a conformational transition between the inactive and active states, leading to a loss of complementarity between the N-SH2 blocking loop and the PTP active site. Previous studies hypothesized that the EF loop of N-SH2, connecting the β-strands E and F, as well as the BG loop, connecting α-helix B and the β-strand G, play an important roles in activation (cf. Figure 1) (27).

**Figure 1.**
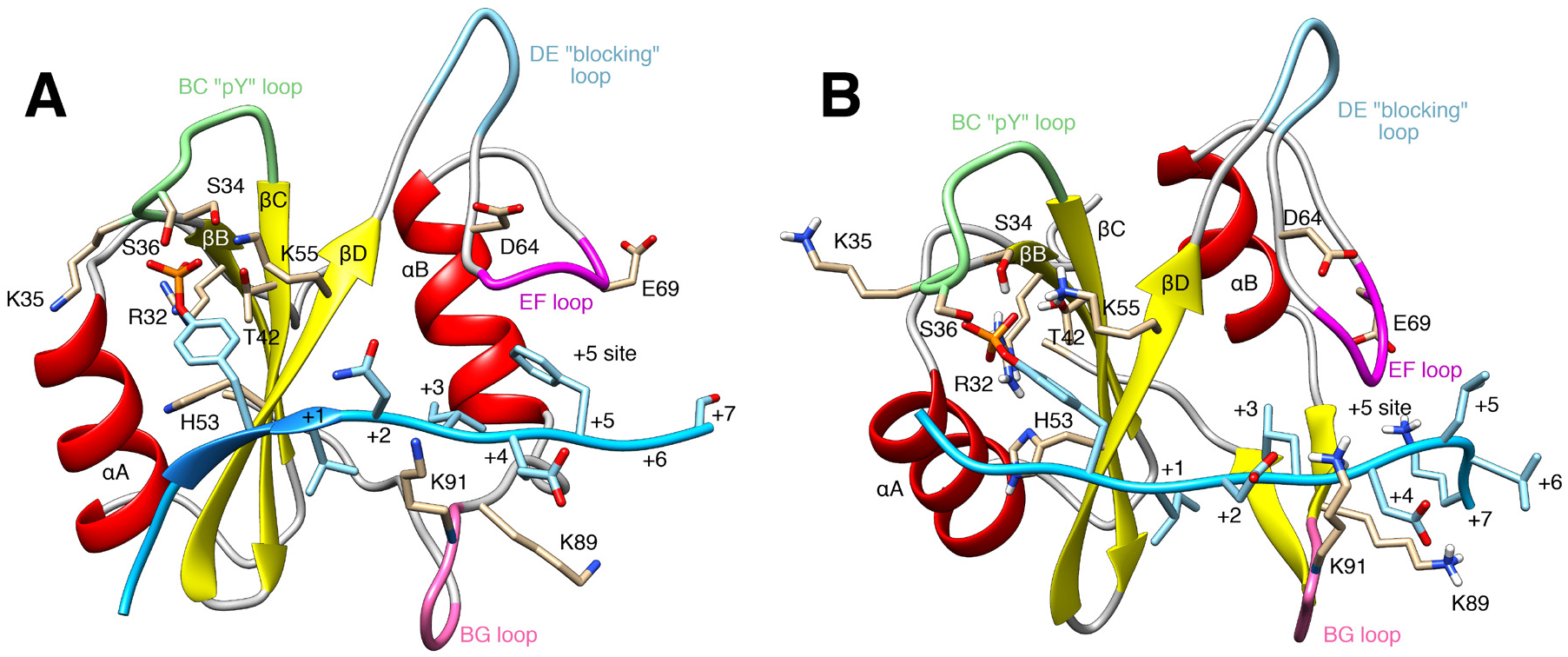
Cartoon representation of the N-SH2 domain in complex with (A) the SPGEpYVNIEFGS peptide [crystal structure (24)] and (B) with the SLNpYIDLDLVK peptide, taken from an MD simulation. The peptide is shown in cyan, comprising the phosphotyrosine and residues from position +1 to +7 relative to the phosphotyrosine (see labels). Functionally important loops are highlighted in color: BC “pY” loop (green), DE “blocking loop” (light blue), EF loop (magenta), BG loop (deep pink). The phosphotyrosine binds the site delimited by the pY loop and the central β-sheet (βB, βC, βD strands). EF and BG loops delimit the “+5 site” where the peptide residue in position +5 is settled.

Two alternative hypotheses have been put forward for the molecular events leading to functional activation of SHP2. First, a “conformational selection” mechanism was proposed based on the observations that *(i)* the crystal structure of the autoinhibited protein shows a completely inaccessible active site, and *(ii)* even under basal conditions in the absence of SH2-binding phosphopeptide ligands, SHP2 exhibits some activity, indicating a transient population of open, active conformations. According to this hypothesis, phosphopeptide binding to the SH2 domains simply stabilizes the fraction of active SHP2, only after dissociation of N-SH2 from the PTP domain has already taken place (7, 17, 28). Second, according to an “induced fit” model, peptide binding might take place in the inactive state of SHP2, triggering the concomitant opening of the protein (29). However, since the N-SH2 binding site is closed in the inactive state, phosphopeptide binding to the inactive state would require a conformational transition of the N-SH2 domain with respect to the crystal structure. A better understanding of SHP2 activation and N-SH2 allostery is the key to clarify how SHP2 function is regulated and how pathogenic mutations cause protein dysfunction.

Guided by extensive molecular dynamics (MD) simulations, various types of free energy calculations, and enhanced-sampling methods, we present an atomistic model of SHP2 activation based on a concerted conformational rearrangement of the N-SH2 domain upon phosphopeptide binding. According the model, N-SH2 predominantly adopts two distinct conformations, denoted α-and β-state, where only the α-state is activating, whereas the β-state stabilizes the N-SH2–PTP interface. Phosphopeptides may bind to the inactive state, subsequently promoting the release of N-SH2 domain from active site; however, the mere phosphopeptide binding is not sufficient for SHP2 activation, as only sequences selecting for the α-state are effective activators. Moreover, the model rationalizes the effect of certain pathogenetic mutations in terms of altered equilibria between the activating and inhibiting N-SH2 conformations.

## RESULTS AND DISCUSSION

Designing an atomistic model of the activation mechanism of SHP2 mediated by the ligand-induced conformational changes requires understanding of the conformational states of the N-SH2 domain. The N-SH2 domain has the two-fold role of blocking the active site of the catalytic domain and recognizing the phosphorylated sequence, the latter triggering the structural rearrangements that precede SHP2 opening and activation.

### Review of N-SH2 structure and phosphopeptide–N-SH2 interactions

Figure 1 presents two representative structures of a phosphopeptide bound to the N-SH2 domain: a crystal structure (24) of the N-SH2 domain in complex with the SPGEpYVNIEFGS peptide (panel A), and a conformation extracted from the MD simulation with the SLNpYIDLDLVK peptide (panel B). Like other SH2 domains, N-SH2 consists of a central antiparallel β-sheet, composed of three β-strands denoted βB, βC and βD, flanked by two α-helices denoted αA and αB. The peptide binds in an extended conformation to the cleft that is perpendicular to the plane of the β-sheet. The phosphotyrosine binding site is delimited by *(i)* the αA helix, *(ii)* the BC loop, hereafter referred as pY loop, connecting two strands of the central β-sheet, respectively βB and βC, and *(iii)* the side chains on the adjacent face of the β-sheet (24).

The key residues involved in the interaction with phosphotyrosine are Arg^32^, Thr^42^, and Lys^55^, lining the central β-sheet, and residues Ser^34^, Lys^35^ and Ser^36^, belonging to the pY loop (24). Residues Ser^34^, Thr^42^ and Ser^36^ form hydrogen bonds with the phosphate group, whereas Arg^32^ and Lys^55^ form salt bridges. Lys^35^ contributes to stabilizing the phosphate group in the binding site by electrostatic interactions. Regarding the peptide sequence, the specificity of peptide–N-SH2 interactions is determined by the peptide residues following immediately after the phosphotyrosine, and interacting with the residues of the N-SH2 domain, placed just on the other side of the central β-sheet (24). Generally, the residues at position +1 and +3 relative to the phosphotyrosine are hydrophobic side chains, pointing towards the N-SH2 domain, while residues at position +2 and +4 are solvent-exposed. Residues at position +2 and +4 interact with the residues belonging to the BG loop, whose sequence represents the principal structural determinant of specificity for the N-SH2 domain. The residue at position +5 is generally inserted into the so-called +5 site, a pocket formed together by the BG and the EF loops. However, also residues further upstream from the phosphotyrosine may interact with N-SH2; for instance, the lysine at positions +7 in the example of Figure 1B favorably interacts with the acidic side chains of residues Asp^64^ and Glu^69^, belonging to the EF loop.

### Phosphopeptides retain the native binding mode throughout the simulations

MD simulations were initially performed on the isolated N-SH2 domain in solution, complexed with a set of phosphopeptides. We considered up to 12 peptides, differing either in length or in sequence and chosen for their high experimental binding affinity or chosen to test the influence of sequence variability (Figure S1). During the simulations, all peptides remained in the binding cleft over the entire 1 μs of simulation, in a conformation very similar to the initial binding mode observed in experimental structures. In Figure S1, the RMS deviation of the most representative MD conformation from the corresponding reference crystal structure is reported for each C_α_ atom of the phosphopeptide. The least-squares fit was performed considering only the domain backbone. Evidently, for all structures the deviation from the experimental binding mode remained below 2-3 Å for positions –1 to +6 relative to the phosphotyrosine, with the exception of GDKQVEpYLDLDLD, where the deviation is below 4 Å. Only the peptide termini exhibit larger RMSD values, indicating weaker structural determinants for binding. Overall, these findings demonstrate that, during the whole simulation, the peptides are tightly bound to the N-SH2 domain. A detailed analysis of ligand–N-SH2 interactions during these simulations are presented elsewhere (30).

### The N-SH2 domain adopts at least two distinct conformations

Whereas the 12 different phosphopeptides maintain their usual binding mode during the simulations, the N-SH2 domain can undergo several conformational transitions. Starting from the crystal structure, in which the pY loop is tightly wrapped around the phosphotyrosine (see Fig. 1A), an opening of the pY loop and the consequent rearrangement of the N-SH2 domain into another distinct structure was observed in several simulations. To characterize the conformations adopted by the N-SH2 domain, principal component analysis (PCA) was applied to the cumulative trajectory of all 12 simulations. Hence, the PCA eigenvectors do not necessarily represent the conformational transitions undergone by the N-SH2 domain during an individual simulation; instead, they describe the overall conformational space accessible to the N-SH2 domain.

PCA suggested that the N-SH2 domain adopts two main conformational states, hereafter called α and β. These two states were resolved by the first PCA vector, which was representative of almost 40% of the overall domain fluctuations. The correlated structural rearrangements detected by the first PCA vector are visualized in Figure 2A by means of the extreme projections onto the vector (see also Fig. 3A), and the related structural rearrangements are quantified in Fig. 2E-G. Namely, the α state shown in Fig. 2A (transparent) is characterized by (i) a closed pY loop (Lys^35^ C_α_–Thr^42^ C_α_ distance ~9 Å), (ii) an increased distance between the ends of two β-strands, βC and βD, leading to breaking of three inter-strand hydrogen bonds and spreading of the central β-sheet into a Y-shaped structure (Gly^39^ C–Asn^58^ N distance ~12 Å), and (iii) the closed +5 site with a narrow, less accessible cleft (Tyr^66^ C_α_–Leu^88^ C_α_ distance ~7 Å; see also Figure 3A, left). The β state shown in Fig. 2A (opaque) is characterized by (i) an open pY loop (Lys^35^ C_α_–Thr^42^ C_α_ distance ~11 Å), (ii) a closed central β-sheet with parallel β strands (Gly^39^ C– Asn^58^ N distance ~4 Å), and (iii) an open +5 site with an accessible cleft (Tyr^66^ C_α_–Leu^88^ C_α_ distance ~12 Å, see also Figure 3A, right).

**Figure 2.**
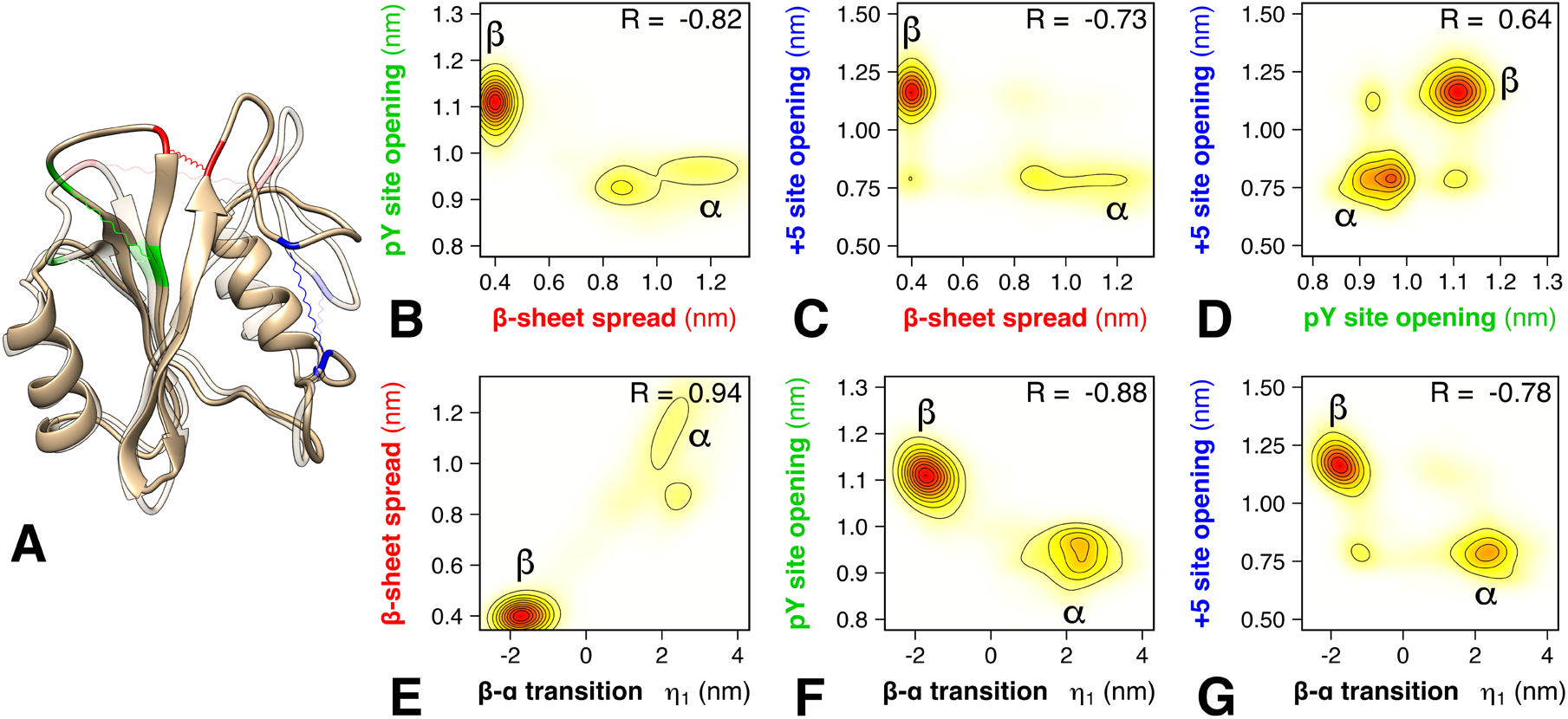
Conformational states and correlations revealed by the first PCA vector. (A) Conformational transition from β (opaque) to α (transparent), representing the two main conformational states adopted by the N-SH2 domain, here visualized as the extreme projections onto the first PCA vector. The residues used to quantify the β-sheet spread, the pY loop opening, and the +5 site opening are highlighted in red, green, and blue, respectively. (B–D) Correlation between the β-sheet spread, pY loop opening, and +5 site opening, as taken from microsecond simulations of N-SH2 bound to 12 different peptides. The distances were defined as described in the main text. (E–G) Correlation between the projection η1 onto the first PCA vector and β-sheet spread, pY loop opening, and +5 site opening (see axis labels). Pearson correlation coefficients *R* are shown in each panel B-G.

**Figure 3.**
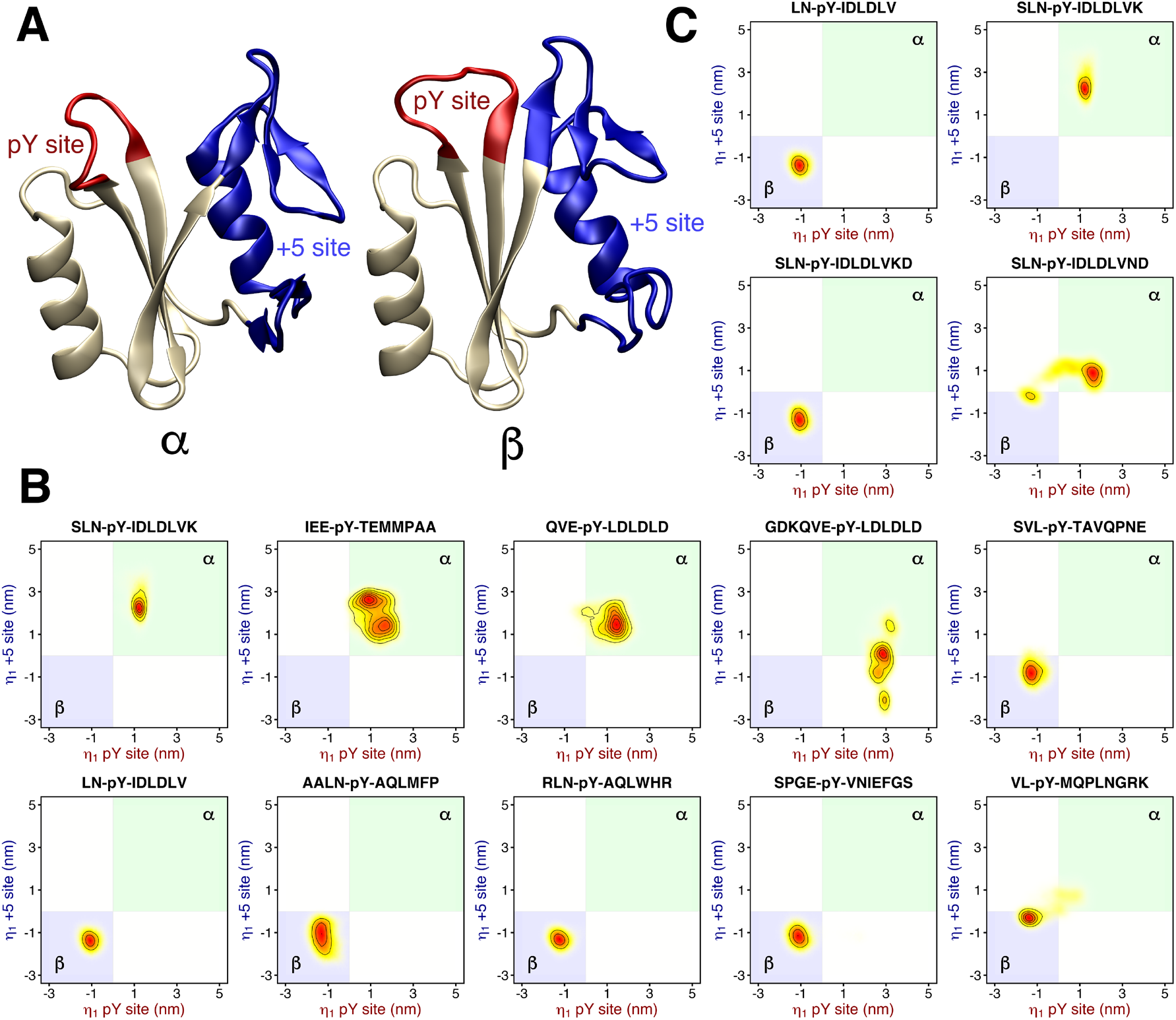
(A) Cartoon representation of the two conformations, α (left) and β (right), adopted by the N-SH2 domain, and shown as the extreme projections onto the first PCA vector. From the first PCA vector, two subsectors were selected, describing the motions of the pY loop (red cartoon) and of the +5 site (blue cartoon), respectively. The residues used for defining the +5 site comprise the end of the βD strand, the DE “blocking” loop, the EF loop, the αB helix and the BG loop. (B/C) Projection of N-SH2 trajectories with different bound peptides (subplot titles) projected onto the PCA subvectors of the pY loop (abscissa) and of the +5 site (ordinate). The region corresponding to the α state (pY loop closed, +5 site closed) is shaded in green, the region of the β state (pY loop open, +5 site open) is shaded in blue.

A correlation analysis showed that the conformation adopted by the pY loop is strictly coupled to the spread of the central β-sheet (Figure 2B, correlation *R* = –0.82). Structurally, this coupling is imposed by a competition between (a) inter-strand hydrogen bonds (one between Gly^39^–Asn^58^ and two between Phe^40^–Ile^56^) of the central β-sheet and (b) hydrogen bonds of Ser^34^ and Ser^36^ of the pY loop with the phosphate group of the phosphotyrosine. Namely, closed inter-strand hydrogen bonds locks the pY loop in an upright, open position, whereas broken inter-strand hydrogen bonds render the pY loop sufficiently flexible for reaching a closed conformation in tight contact with the phosphotyrosine.

The correlations between the spread of the central β-sheet and pY loop (Fig. 2C, *R* = –0.73) as well as between the pY loop and the opening/closure +5 site (Fig. 2D, *R* = 0.64) are slightly weaker yet clearly detectable. The latter correlation represents a potential mechanism for modulating the conformational rearrangement of the N-SH2 domain, through the recognition of a particular peptide sequence at the +5 site, as analyzed in the following in detail.

### The conformational changes of pY loop and of the +5 site are highly concerted

Our simulations suggest a coupling between the pY loop and the +5 site, revealing an allosteric interaction for controlling the state of the pY loop through binding of a specific amino acid sequences at the +5 site (Fig. 2D). To analyze the coupling during simulations with different peptides, we split the PCA vector into two subvectors that describe the motion either of the pY loop or of the +5 site. In principle, projections on these two subvectors could characterize four conformational states given by the two possible states of the pY loop times two possible states of the +5 site (four shaded areas in each panels of Figure 3B/C). The two additional states, relative to the α and β states characterized above, would be given by (i) a closed pY loop with an open +5 site and (ii) an open pY loop with a closed +5 site (Fig. 3B/C, white areas in panels).

As reported in Figure 3B/C, a correlation between the pY site and the +5 site is observed in all simulations, where most peptides strictly select for either the α or for the β state (Fig. 3B/C, blue/green areas). In other words, when the pY site is closed, the +5 site is mostly closed (β state); when the pY site is open, the +5 site is always open (α state). Consequently, a conformational change at the pY site is accompanied by concerted conformational change at the +5 site, spanning ~20 Å across the N-SH2 domain. These results support the presence of an allosteric mechanism that couples two binding sites with different physicochemical properties: the pY site, characterized by a high affinity for phosphotyrosine but lacking specificity, and the +5 site, whose role is to induce specificity in the binding of different amino acid sequences.

### The phosphopeptide binding mode and sequence play important roles in selecting the N-SH2 domain conformation

The equilibrium MD simulations showed that each peptide may induce the N-SH2 domain to populate mostly one among the two conformations, and the simulations revealed a correlation between the pY and +5 sites. However, it is unlikely that each individual simulation may explore all possible binding modes of a phosphopeptide within 1 μs of simulation time. Indeed, the interconversion between binding modes would first involve a weakening of peptide–N-SH2 interactions, before the required side chain and backbone reorientations may occur. Consequently, the equilibrium MD simulations were insufficient for unambiguously deciding whether a specific peptide sequence selects for the α or the β state.

However, the simulations did reveal how specific binding modes may lock N-SH2 in the α or β state. Figure 3C compares the simulation of the SLNpYIDLDLVK peptide with simulations of three peptide analogues with modifications at positions +6, +7 and +8, thereby adopting different polarities at the capped carboxy terminus. These subtle differences in the sequence affected the orientation of the residue in position +5. In case of SLNpYIDLDLVK and SLNpYIDLDLVND, the leucine side chain at position +5 was exposed to the solvent, thereby allowing closure of the +5 site and consequently the population of the α state. In contrast, in case of SLNpYIDLDLV and SLNpYIDLDLVKD, the leucine side chain points towards the binding cleft, keeping the +5 site open and the N-SH2 domain in the β state.

To investigate in more detail the role of the residue at the +5 site in imposing the N-SH2 conformation, we carried out a large set of additional free energy calculations using non-equilibrium transitions and Crooks Fluctuation Theorem (31, 32). To this end, we computed the change of preference of the ligand/N-SH2 complex for the α or the β state upon introducing a mutation in the ligand, according the thermodynamic cycle reported in Figure S2. Figure S3 presents the computed ∆∆*G* values for the ligands SLNpYIDLDL+5VK and SPGEpYVNIEF+5GS for all possible mutations (except proline) at position +5 and, as a control, at position +6. Positive ΔΔ*G* indicates an augmented preference for the α state, and a negative ΔΔ*G* an augmented preference for the β state. Evidently, mutations at the +5 position modulate the α-versus-β preference by up to 14 kJ/mol, whereas mutations at the +6 position have a much smaller effect. Generally, bulky hydrophobic aminoacids at +5 such as Phe, Leu, Ile seem to stabilize the β state (Figure S4), whereas substitutions with bulky polar amino acids such as Asp, Glu, Arg, Asn, or Gln seem to favor the α state (Figure S5), although exceptions exist. These trends in stabilizing α versus β, as derived from the free energy calculations, are well confirmed by additional free simulations (Figure S4 and S5).

Taken together, these data further corroborate that the N-SH2 domain may adopt two alternative conformations, constituted by a coupling between *(i)* the pY loop, which binds the phosphotyrosine, and *(ii)* the +5 site, which recognizes the peptide sequence downstream of the phosphotyrosine. However, the preference for one of the two alternative conformations, α or β, depends not only on the particular peptide sequence but also on the binding mode adopted by the ligand. In particular, both the polarity and the spatial orientation of peptide residue at the +5 site emerge as key determinants for the N-SH2 conformation.

### Free-energy calculations confirm that the ligand residue at the +5 site is a key determinant for the preference N-SH2 for the α or β state

To overcome sampling problems of unbiased MD simulations and to rationalize the preference for the α or β states in energetic terms, we computed the free energy profile of the α– β conformational transition using umbrella sampling. Here, we used the distance between the central β-strands as the reaction coordinate because *(i)* this distance is strongly correlated with the α–β transition (see Figs. 2B, C, E) *(ii)* pulling along a center-of-mass distance may be less prone to sampling problems as compared to pulling along a collective PCA vector.

The conformational equilibrium between the α and β states is determined by a balance of different energetic contributions: *(i)* the intermolecular interactions between the ligand and the N-SH2 domain, and *(ii)* the intramolecular interactions between the residues of N-SH2, as rationalized for three typical binding modes:

- In absence of a ligand (Figure 4A), the β state represents the free-energy minimum, presumably favored by the formation of inter-strand H-bonds between the closed central β-sheet. In turn, increased free energy is required for spreading the β-strands into a Y-shaped structure, so that adopting the α state is unfavorable.
- In presence of a truncated ligand, such as SPGEpYVNI (Figure 4B), which contains a phosphotyrosine but lacks the residues flanking the +5 site, the α state is the free-energy minimum. In this case, the pY loop–phosphate interactions prevail over inter-strand H-bonds of the central β-sheets, leading to closure of the pY loop and opening of the β-sheets.
- With the ligand SPGEpYVNIEFGS, the β state is the free-energy minimum (Figure 4C). Here, a bulky phenylalanine at position +5 occupies the cleft of the +5 site, thereby keeping the +5 site open. This conformation stabilizes the central β-sheet, aiding the inter-strand H-bonds of the β-sheet to prevail over the pY loop–phosphate H-bonds. Consequently, β-sheets close and pY-loop opens.

**Figure 4.**
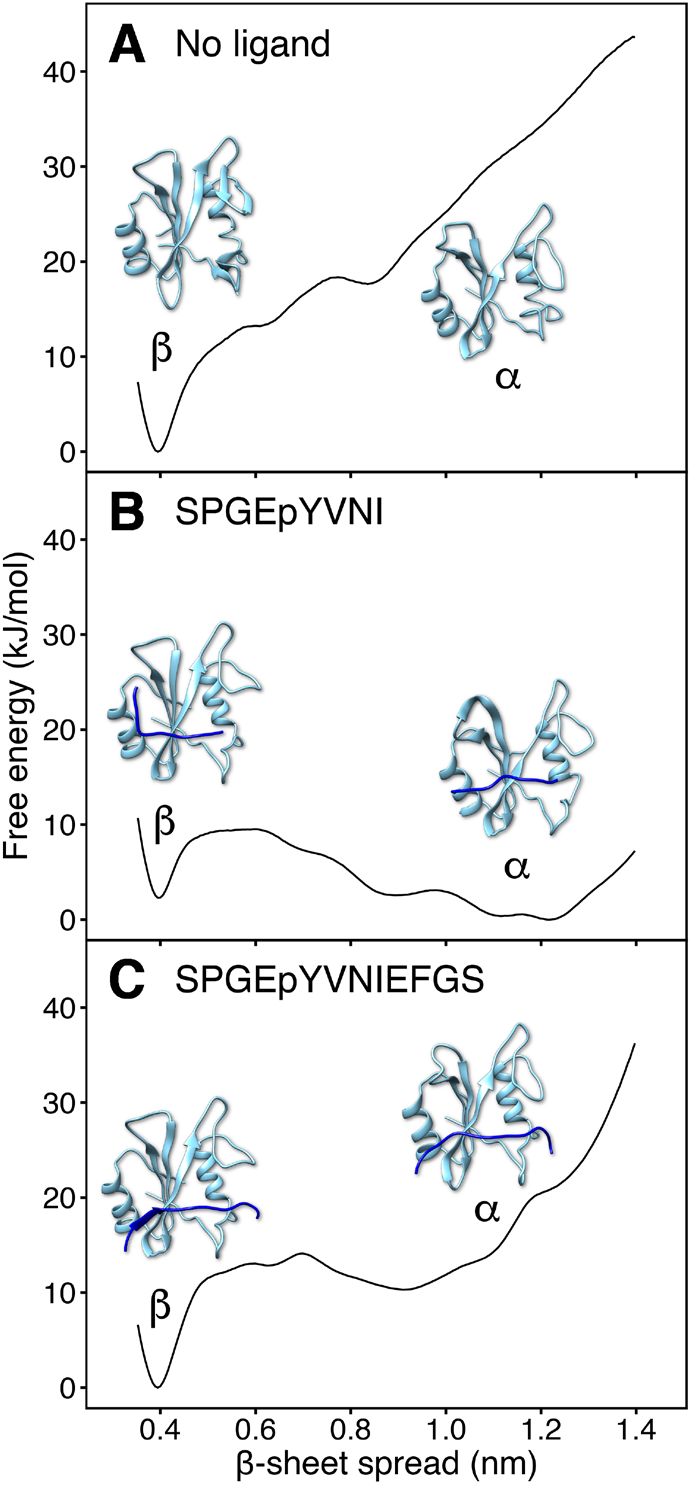
Free energy profiles of the opening of the central β-sheet for the N-SH2 domain in apo state (no ligand), bound to a truncated peptide (SPGEpYVNI), or bound to a full-length peptide (SPGEpYVNIEFGS). The opening of the central β-sheet was quantified as the distance between the backbone N and C atoms of residues Gly^39^ and Asn^58^, respectively.

The free energy profiles confirm that a subtle balance between *(i)* ligand/N-SH2 interactions and *(ii)* intradomain interactions within N-SH2 determine the conformation of N-SH2. The ligand residue at the +5 position plays a key role in imposing the α versus the β state. However, because the side chain at ligand position +5 may adopt different orientations relative to the +5 site (towards the cleft or solvent-exposed), not only the type of side chain, but also the binding mode of the overall ligand may influence the N-SH2 conformation.

### The N-SH2 domain populates the β state in the autoinhibited state of SHP2

One of the main features that distinguishes the α state from β, namely the spread of the central β-strands, is also revealed by comparing the crystallographic structures of the autoinhibited conformation of SHP2 with those of the N-SH2 domain bound to a phosphopeptide. Table S1 reports the distance between the two central β-strands, βC and βD, for each crystal structure. Evidently, in the autoinhibited conformation of SHP2, the distance between the β-strands is small (3.7 to 4.0 Å) indicative of closed β-sheets (23, 26). Hence, N-SH2 bound to PTP adopts a conformation similar to the β state observed in simulations. In contrast, when N-SH2 is complexed with a phosphopeptide, this distance is greater (~7-8 Å) indicative of open β-sheets, a peculiarity of the α state (24).

Whereas the central β-strands of N-SH2 have clearly taken the β-state in crystal structures of autoinhibited SHP2, as reported in the previous paragraph, the structures of the pY loop and of the +5 site have been crystallographically less well defined. The pY loop was typically modeled with a partly open conformation, corresponding to an intermediate conformation between the α-and β-states described here (2SHP, 4DGP) (23, 26). However, the electron densities at the pY loop were much lower compared to the density of the nearby β-strands (e.g. 2SHP, 4DGP, 4GWF) (23, 26), indicating increased disorder of the pY loop. Further, the partially closed pY loop might have been stabilized by crystal contacts (e.g. 2SHP, 4DGP) (23, 26) and by a buffer phosphate bound to the phosphotyrosine binding site, as modeled by the 5EHP structure (33). Indeed, during the MD simulations with CHARMM36m or different Amber force fields, the pY loop of autoinhibited SHP2 reproducibly opened up, taking the open pY loop conformation of the β-state. Likewise, the electron density of the BG loop, which forms the pocket of the +5 site together with the EF loop, was poorly defined in several SHP2 crystals and, consequently, only partly modeled against the data by several authors (5EHP, 4GWF) (33). Again, during free simulations of apo SHP2 in solution, we repeatedly observed a complete opening of the cleft at the +5 site, consistent with the β state. Taken together, crystallographic data unambiguously showed that that central β-strands of N-SH2 take the β-state in the autoinhibited state of SHP2. For the pY and +5 site, crystallographic data is more ambiguous and, hence, simulations presented here were needed to reveal that also the pY and +5 site take the β-state in autoinhibited SHP2.

### The α state promotes SHP2 opening and activation

Based on the observation that the β state is found in association with the autoinhibited SHP2, whereas in crystal structures, the isolated, peptide-bound N-SH2 structure resembles the α state, we hypothesized that the α state represents an activating state. According to this model, binding of an activating phosphopeptide would trigger the first steps of SHP2 activation by switching the N-SH2 domain towards the α conformation and, subsequently, weakening the interactions between the blocking loop and the catalytic site.

To test this model, we used umbrella sampling to compute the free energy profiles of SHP2 activation with the N-SH2 domain restrained either to the α or to the β state. Here, the restraint to α or β was implemented by restraining a single principal component, while leaving all other degrees of freedom unbiased, hence representing only a mild restraint (cf. Methods). As a reaction coordinate for SHP2 activation, we used the center-of-mass distance between the backbone atoms of blocking loop and the catalytic PTP loop. Upon pulling the simulation system along this coordinate, the N-SH2 domain moved from its position in the autoinhibited state (Figure 5A) to a different position on the PTP surface (Figure 5B), typically by sliding over the PTP surface. The final position of N-SH2 differed among independent simulation runs, indicative of a large accessible conformational space of activated SHP2.

**Figure 5.**
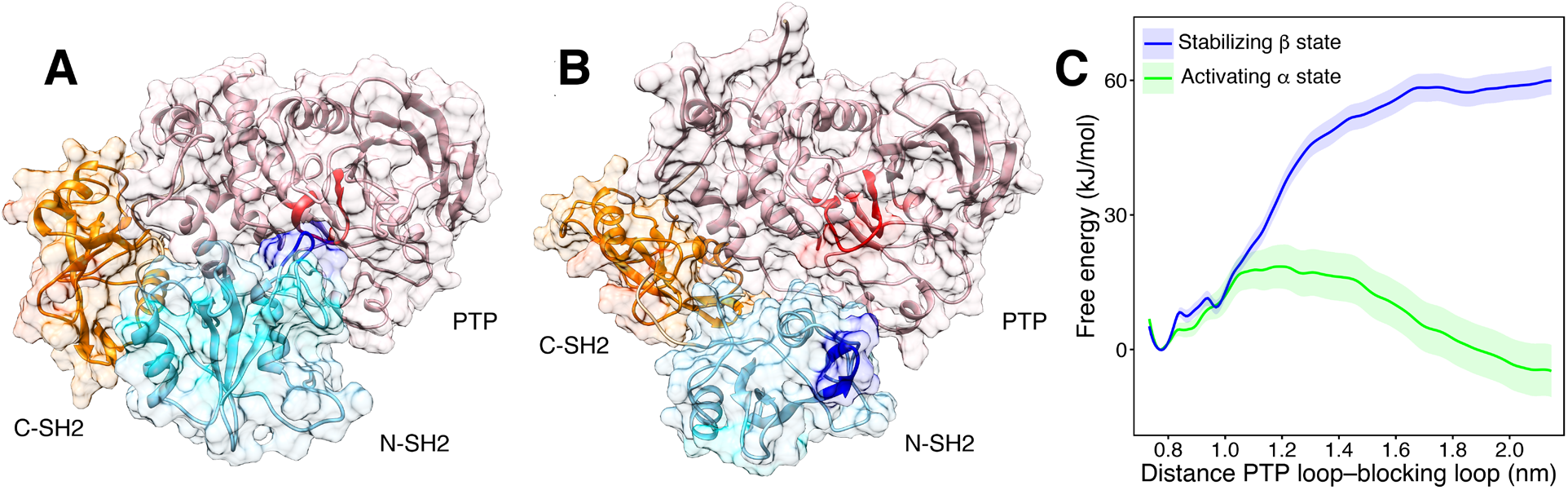
(A) In the autoinhibited structure of SHP2, the N-SH2 domain (cyan cartoon) blocks the catalytic site (red) of the PTP domain (pink) with the blocking loop (blue). The N-SH2 domain is connected to PTP in tandem with the homologous C-SH2 domain (orange). (B) Open and active structure obtained by pulling simulation coupled with simulated tempering, using the color code of panel A. (C) Free energy profiles of SHP2 opening with the N-SH2 restrained to the α state (green line) or to the β state (blue line). The free energy calculations were performed using CHARMM36m force field. The distance between the blocking loop and the catalytic PTP loop, in terms of distance between the backbone centers-of-mass of residues 60–62 and residues 460–462, was taken as reaction coordinate.

In agreement with the model, we found that the N-SH2 state strongly influences the free energy of activation (Figure 5C). Namely, according to the free energy profile for the N-SH2 domain restrained to the β state, a high free energy penalty is associated with SHP2 activation, indicating that the β state favors a stable autoinhibited SHP2 conformation (Figure 5C, blue line). In contrast, upon restraining N-SH2 in the α state, SHP2 activation is associated with only a small barrier, and the active state represented the free-energy minimum (Figure 5C, green line). The exclude that the choice of the reaction coordinate influences these findings, we computed a second set of free energy profiles along the distance between the C_α_ atoms of residues Asp^61^ and Ala^461^, representing central residues of the blocking loop and catalytic PTP loop, respectively. These profiles were qualitatively similar to the first set of profiles (Figure S6). Taken together, with both the reaction coordinates, we consistently obtained greatly increased free energy for activation in the β state as compared to the α state, corroborating that the α state destabilizes the N-SH2/PTP interface, thereby promoting activation.

### Amino acid substitutions at codon 42: basal activity and response to ligand stimulation are rationalized with altered conformational selection between the α and β state

Many pathogenic mutations of SHP2 cluster at the N-SH2/PTP interface, where they may destabilize the N-SH2/PTP interactions, causing constitutive activation of SHP2 (29). However, certain pathogenic mutations seem to have more subtle, allosteric effects, since their impact on SHP2 function cannot be rationalized by mere steric effects. Here, we study such mutations to test our proposed model of SHP2 activation. A typical example is the Noonan Syndrome (NS)-causing Thr42Ala substitution that replaces a conserved threonine in the central β-sheet. Because Thr^42^ forms an H-bond with the phosphotyrosine in wild-type (wt) N-SH2, one might expect that Thr^42^ contributes to the stability of the N-SH2/phosphopeptide complex. However the substitution with alanine leads to an increase in phosphopeptide binding affinity (6), as documented by the dramatically enhanced catalytic activity of the SHP2^A42^ mutant following stimulation with a bisphosphoryl tyrosine-based activation motif (BTAM) peptide (6, 28).

Notably, among all possible mutants arising from single base changes in codon 42, all but two SHP2 mutants exhibit a basal catalytic activity comparable to wild-type SHP2. The exceptions are SHP2^P42^ that showed a threefold increase, and SHP2^I42^ that showed a 50% increase (6). Upon BTAM peptide stimulation, mutants not associated with NS were either responsive (SHP2^S42^, SHP2^R42^), with a variable increase of the catalytic activity respect to the basal condition, or unresponsive (SHP2^P42^, SHP2^I42^, SHP2^K42^). Relative to the wild-type protein, SHP2^S42^ exhibits a 50% higher stimulated activity, whereas in SHP2^R42^ the stimulated activity was significantly lower (6). Surprisingly, despite the different range of catalytic activities exhibited by these mutants, the binding affinities were only moderately affected, with the exception of the above-mentioned SHP2^A42^(6).

These modifications of basal activity and responsiveness to peptide stimulation cannot be easily explained by perturbed interaction between residue 42 and the phosphotyrosine. Previous simulations, which showed reduced fluctuations of the pY loop in the Thr42Ala mutant did not reveal a molecular mechanism of the mutation effects (6). Therefore, we hypothesized that the overall change of the catalytic activity might derive from a combination of two mechanisms: *(i)* the perturbation of phosphotyrosine interaction upon amino acid substitution, which may affect both the α and the β state, *(ii)* the shift of the equilibrium between the activating α state and the stabilizing β state, which may arise from the replacement of the conserved threonine at the central β-sheet.

We computed the free energy profile for the opening of the central β-sheet, as required for the β-to-α transition, for wild-type N-SH2 and for various mutants of the domain in absence of phosphopeptide (Figure 6A/B). Remarkably, the Thr42Ile mutant greatly stabilized an open β-sheet conformation corresponding to the activating α state, which is compatible with the experimentally observed increased basal activity of SHP2^I42^(Figure 6B, green solid line) (6). All other simulated mutants had a smaller effect on the conformation of the central β-sheet, in line with a basal activity of those mutants similar to the wild type (Figure 6B). These findings support the hypothesis that mutations may modify the basal activity of SHP2 by shifting the conformational equilibrium of apo N-SH2. In addition, the simulations corroborate our hypothesis that the α state, as stabilized by the Thr42Ile mutant, is indeed an activating state.

**Figure 6.**
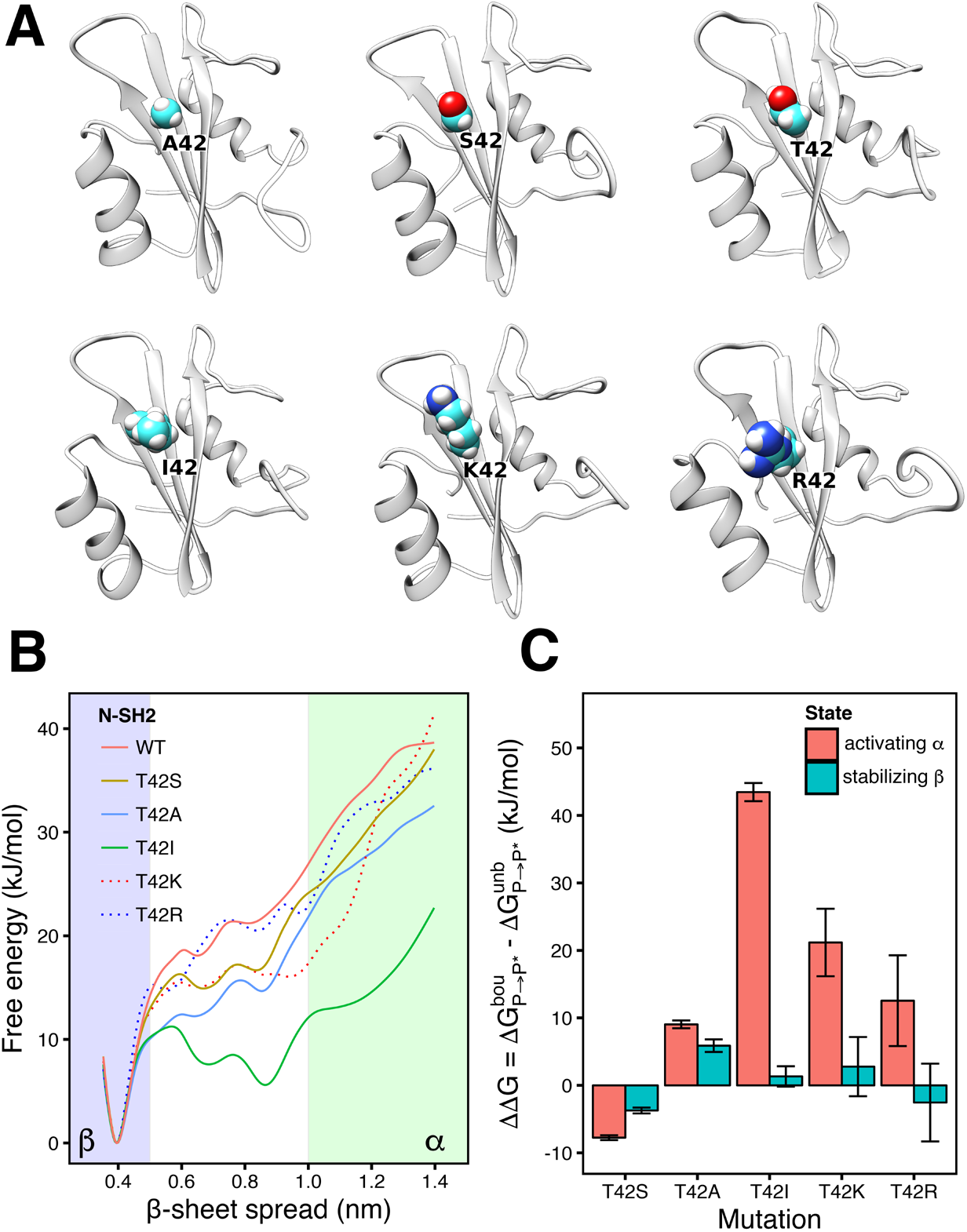
(A) Cartoon representation of N-SH2, wild type and Thr^42^ mutants (A42, S42, I42, K42, R42), sorted according the size of the sidechain (Ala < Ser < Thr < Ile < Lys < Arg). (B) Free energy profile for the opening of the central β-sheet in wild type and T42 mutants, as a function of the N–C distance between the residues Gly^39^ and Asn^58^. Statistical errors were <2 kJ/mol (not shown for clarity). (C) ΔΔG of binding of the T42 mutants respect to the wild-type form, calculated for the N-SH2 domain in α or in β state.

Next, we evaluated whether mutations at codon 42 may modulate the binding affinity of a peptide to N-SH2 either in the activation α state or in the stabilizing β state. Because it is difficult to sample a complete ligand binding process or the α–β transition in the presence of a peptide, we used thermodynamic cycles and alchemical free-energy calculations based on Crooks Theorem (31, 32). Accordingly, we here computed the free energies for alchemically transforming Thr^42^ to amino acids Ser, Ala, Ile, Lys, or Arg, either in presence (denoted Δ G_P-P_ * [bou]) or absence (Δ G_P-P_ * [unb]) of a bound reference peptide SLNpYIDLDLVK. Using a thermodynamic cycle, the difference between these two Δ G-values, denoted ΔΔG, yields the change in peptide binding affinity upon introducing the mutation. Critically, because these calculations were carried out with N-SH2 restrained either in the α or in the β state, the computed ΔΔG(α) and ΔΔG(β) values show whether a Thr^42^ mutation (de)stabilizes ligand binding to the α state, to the β state, or to both.

We found that the Thr42Ile, Thr42Lys, Thr42Arg substitutions greatly affect the affinity of the peptide to the activating α state, as evident from large positive ΔΔG values in a range of tens of kilojoule per mole (Figure 6C, red bars). However, the same mutations only moderately affect the peptide affinity to the β state, as shown by ΔΔG the range of only a few kilojoules per mole (Figure 6C, blue bars). Hence, upon introducing these amino acid changes, the peptide may still bind to SHP2, but N-SH2 gets locked in the β state thereby stabilizing the autoinhibited state of SHP2. In turn, the activating α state may be greatly destabilized, thereby rendering the mutations Thr42Ile and Thr42Lys completely unresponsive to ligand stimulation (ΔΔG(α) values of +43kJ/mol or +21kJ/mol, Fig. 6C), in qualitative agreement with the experimental findings (6). The substitution Thr42Arg with a ΔΔG(α) value of +13kJ/mol only moderately destabilizes the α state, thereby rendering the mutant less responsive than the wild type, qualitatively in line with experimental data (6).

Structurally, the perturbations of the affinity to the α and β states in Thr42Ile, Thr42Lys and Thr42Arg SHP2 mutants may be rationalized considering that any substitution to a larger side chain impedes the formation of the directed hydrogen bonds involved in the closed pY loop. In contrast, in the β state, the substitution with isoleucine does not significantly affect the salt bridges with domain residues Arg^32^ and Lys^55^, and the steric hindrance of arginine and lysine is balanced by the new electrostatic interaction with the phosphate group.

The ΔΔG values for the Thr42Ser and Thr42Ala substitutions are qualitatively different compared to the Thr42Ile, Thr42Lys and Thr42Arg (Fig. 6C). For Thr42Ser and Thr42Ala, the respective ΔΔG(α) and ΔΔG(β) are similar, suggesting that ligand binding does not lock N-SH2 in the stabilizing β state but instead ligand binding still allows population of the activating α state. Consequently, Thr42Ser and Thr42Ala are responsive to ligand stimulation, again in qualitative agreement with the experiment (6). Taken together, both the basal activity of mutants at codon 42 as well as their response to ligand stimulation correlate with modified α–β equilibria revealed by the free-energy calculations, corroboration that α is an activating state.

### The increased peptide affinity of Thr42Ala might be associated with a modified α–β equilibrium

In line with the available experimental observations, Thr42Ile, Thr42Lys and Thr42Arg have only a small effect on the overall ligand affinity, as evident from ΔΔG(β) values close to zero; instead, these mutations merely shift the equilibrium of the peptide-bound N-SH2 towards the β state (see above). Rationalizing experimental ligand affinities of Thr42Ala and Thr42Ser is, however, more difficult in the light of the ΔΔG values. Namely, for the Thr42Ala N-SH2, we found slightly positive ΔΔG values for both the α and the β state, indicating a reduced ligand affinity (Figure 6C). For Thr42Ser, we found slightly negative ΔΔG values, indicating an increased ligand affinity. These ΔΔG values are structurally expected because Thr42Ala leads to a loss of the H-bond of residue 42 with the phosphotyrosine, whereas this H-bond is maintained in Thr42Ser. However, at first sight, these values seem to contradict surface plasmon resonance experiments that reported an increased ligand affinity to Thr42Ala and an unmodified affinity to Thr42Ser (6).

We hypothesize that the structurally unexpected, but experimentally found, increased affinity to Thr42Ala may be rationalized by a modified α–β equilibrium of apo N-SH2. The free energy profiles for the opening of the central β-sheet suggest that a substitution of Thr42 with an aliphatic residue, such as alanine or isoleucine, renders the N-SH2 domain more prone than the wild type to spread the β-strands into a Y-shaped structure and, hence, to adopt the α state (Figure 6B, compare cyan/green lines with red line). We expect that such increased population of the α state would facilitate peptide binding because, only in the α state, the pY loop is tightly wrapped around the phosphotyrosine, thereby forming strong polar interactions with the peptide.

To validate this model of Thr42Ala affinity in a quantitative manner, calculations of the absolute peptide binding affinity, possibly with N-SH2 restrained to the α or to the β state, as well as structural studies will be required in the future. However, considering that modulations of conformational equilibria are widely adopted by enzymes to control ligand affinity (34), a modified α–β equilibrium is a plausible mechanism of the increased peptide affinity of Thr42Ala.

## CONCLUSIONS

We presented an atomistic mechanism for activation of SHP2, mediated by ligand-induced conformational changes of the N-SH2 domain. Extensive simulations showed that N-SH2 in solution adopts two distinct conformations, denoted α and β. The α and β states exhibit different structures of the central β-sheet and of two ligand interaction sites, which are responsible respectively for the binding of the phosphotyrosine (pY site) and for recognizing peptide residues downstream of the phosphotyrosine (+5 site).

The simulations revealed an allosteric interaction between the pY and +5 sites, mediated by the conformation of the central β-sheet: when the pY site is closed, the +5 site is mostly closed, with the central β-sheet arranging in a Y-shaped conformation; when the pY site is open, the +5 site is always open, with the central β-sheet retaining parallel β-strands.

In absence of a ligand, the N-SH2 domain remains in the β state, both in its isolated form and in the autoinhibited structure of SHP2. Peptide binding may either lock the N-SH2 domain in the β state or trigger the transition to the α state, depending on the sequence and the binding mode of the peptide around the +5 site. Hence, the allosteric interaction provides a means for controlling the conformations of both the pY site and of the central β-sheet by specific peptide sequences and peptide binding modes at the +5 site.

A ligand-induced transition from the β to the α state has two consequences; *(i)* the phosphotyrosine is bound more tightly because only in the α state the pY loop is sufficiently flexible in order to fully close onto the phosphate group, thereby forming stronger polar interactions. This mechanism provides a means for controlling ligand affinity via the conformation at the +5 site; *(ii)* in SHP2, with N-SH2 in the α state with a Y-shaped central β-sheet, interactions between the N-SH2 domain and the catalytic PTP domain are weakened, thereby facilitating the dissociation of N-SH2 and hence the activation of SHP2. According to this model, peptide binding to N-SH2 is necessary but not sufficient for SHP2 activation as only those peptides that select for the α state are activating (Figure 7).

**Figure 7.**
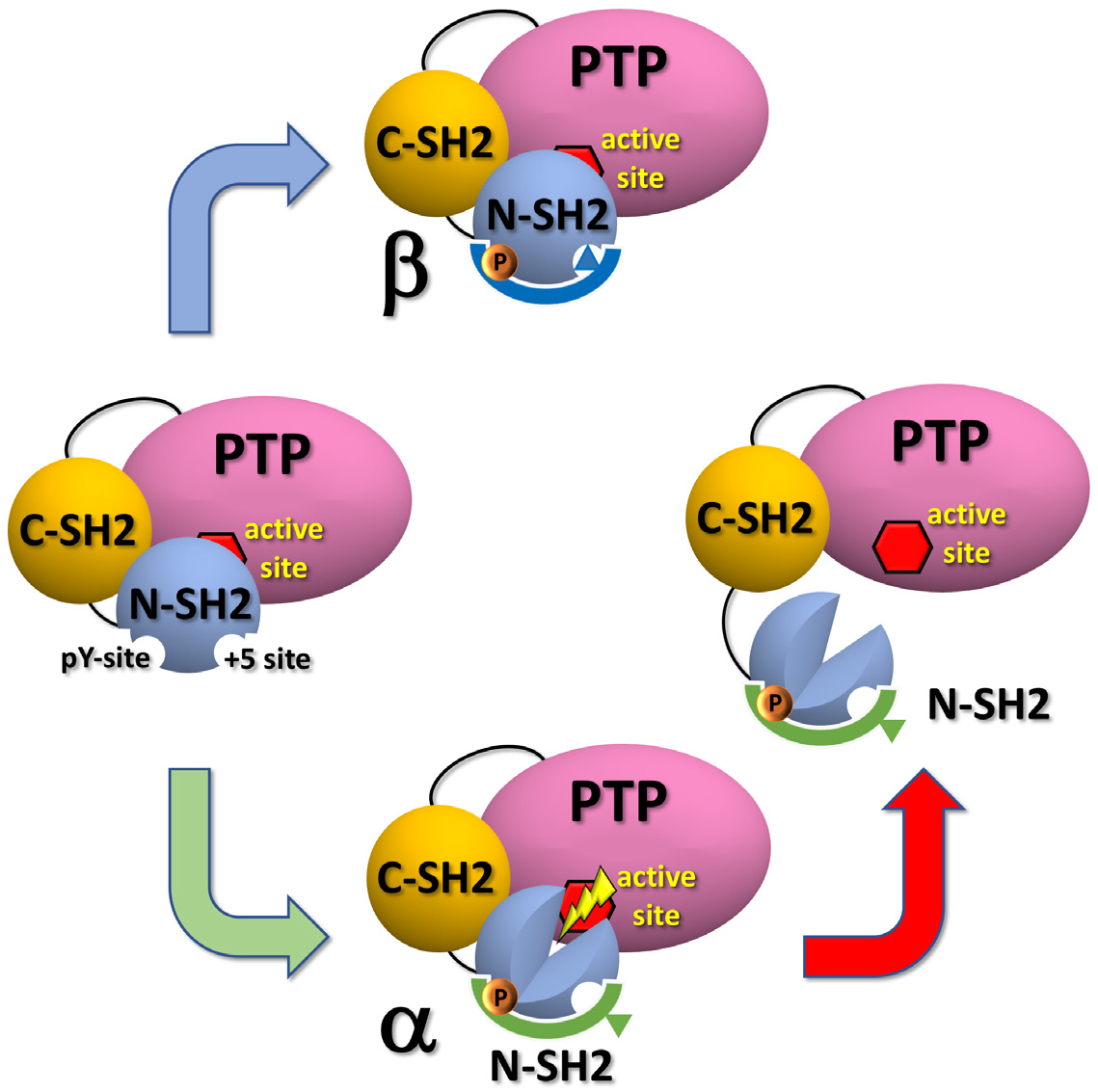
Simplified scheme of SHP2 activation, triggered by a ligand binding-induced conformational transition of the N-SH2 domain. In the apo form, in absence of a peptide, N-SH2 adopts the β state, leaving SHP2 mostly a closed autoinhibited structure (left panel). Binding of certain peptides may lock N-SH2 in the β state, stabilizing the autoinhibited conformation of SHP2 (upper panel). SHP2 activation proceeds only if the peptide promotes the transition of the N-SH2 domain towards the α state, leading to a Y-shaped central β-sheet, thereby weakening the N-SH2–PTP interactions (lower panel). N-SH2 dissociates from PTP, thereby exposing the active site of PTP to the solvent (red hexagon, right panel).

The model of SHP2 activation was further validated by rationalizing the effects of amino acid substitutions of Thr^42^ in terms of altered equilibria between the α and the β state. We found that Thr^42^ mutations may shift the α–β equilibrium of N-SH2 both in the apo form and in contact with ligands, thereby modulating the basal activity (Thr42Ile) or the responsiveness to ligand stimulation, in qualitative agreement with experimental data. We further hypothesized that an altered α–β equilibrium may underlie the structurally unexpected increased peptide affinity of the Thr42Ala mutant.

Understanding of the activation mechanism of SHP2 also forms the basis to a mechanistic understanding of SHP2 inhibition by small molecules or peptide mimics and, consequently, such understanding may guide the development of new therapeutic options. Specifically, our finding that the selection of particular binding modes leads to the activation, rather than mere binding of the effector, opens novel strategies for SHP2 regulation. As such, we hope that the allosteric activation mechanism of SHP2 proposed here will support ongoing efforts against genetic disorders involving SHP2 (35, 36, 37, 38).

## METHODS

### MD simulations of the N-SH2 domain complexed with different phosphopeptides

Molecular dynamics simulations were performed on the isolated N-SH2 domain in solution, complexed with a set of phosphopeptides, comprising SLNpYIDLDLVK, IEEpYTEMMPAA, QVEpYLDLDLD, SVLpYTAVQPNE, AALNpYAQLMFP, RLNpYAQLWHR, SPGEpYVNIEFGS, VLpYMQPLNGRK and their analogs. The initial atomic coordinates were derived from crystallographic structures (Table S2). For some peptides (e.g., SPGEpYVNIEFGS) the sequence in the simulation exactly matched the original sequence, and the crystal structure has been used without any modification. For other peptides (e.g., SLNpYIDLDLVK), the crystal structure with the highest peptide sequence similarity was chosen, and the structure was edited by means of Molecular Operative Environment (MOE) (39), followed by a conformational analysis and a local energy minimization with side chain repacking, yielding a reasonable binding pose for all peptides (Table S2). The termini of peptides were capped by acetyl and amide groups. N-SH2 domains in simulation comprised the residues from position 3 to 103. Each protein molecule was put at the center of a dodecahedron box, large enough to contain the domain and at least 0.9 nm of solvent on all sides. Thus depending the specific length and charge of the peptides, the protein was solvated with ~4400–6700 explicit TIP3P water molecules (40), and up to 6 Na^+^ ions were added to neutralize the system. All MD simulations were performed with the GROMACS software package (41), using AMBER99SB force field if not stated otherwise (42), augmented with parm99 data set for phosphotyrosine (43). Long range electrostatic interactions were calculated with the particle-mesh Ewald (PME) approach (44). A cutoff of 1.2 nm was applied to the direct-space Coulomb and Lennard-Jones interactions. Bond lengths and angles of water molecules were constrained with the SETTLE algorithm (45), and all other bonds were constrained with LINCS (46). The pressure was set to 1 bar using the weak-coupling barostat (47). The temperature was controlled at 300 K using velocity rescaling with a stochastic term (48). For all systems, the solvent was relaxed by energy minimization followed by 100 ps of MD at 300 K, while restraining protein coordinates with a harmonic potential. The systems were then minimized without restraints and their temperature thermalized to 300 K in 10 ns, in a stepwise manner. Finally, production simulations of 1 μs were performed.

### Principal component analysis (PCA) of the N-SH2 domain dynamics

PCA was performed on a cumulative trajectory of all simulations of the N-SH2 domain, after removing ligand coordinates. The structures were superimposed by a least-squares fit on the backbone considering only the residues with small root mean-squared fluctuation, representing the relatively rigid core of the domain. The core was defined by the sequence ranges Phe^7^-Pro^33^, Asp^40^-Arg^47^, Ala^50^-Asn^58^, Asp^61^-Leu^65^, Phe^71^-Tyr^81^, Leu^88^-Glu^90^, Val^95^-Pro^101^. This procedure avoids that the fits are influenced by large fluctuations of loops and termini. The covariance matrix was built using backbone atoms of residues Trp^6^ to Pro^101^, excluding the flexible termini.

### MD simulations of the activation of SHP2, with N-SH2 restrained into the α or β state

The initial coordinates of SHP2 were taken from the autoinhibited conformation (2SHP) (23). The protein was positioned at the center of a dodecahedral box, large enough to contain the protein and at least 0.9 nm of solvent on all sides, and solvated with ~23,000 explicit water molecules and three Na^+^ ions. Because certain Amber force fields in conjunction with TIP3P have been reported to overstabilize protein-protein contacts (49), we used the CHARMM36m force field for simulations of SHP2 activation (50). The structure was minimized and thermalized to 300 K using the procedure described above. A simulation of 1 μs was performed to ensure a well equilibrated autoinhibited structure in solution. Next, the N-SH2 domain of SHP2 was restrained in one of two distinct states (denoted as α-and β-state) using a harmonic restraint on a single principal component of C_α_ atoms from residue Trp^6^ to Pro^101^(k = 1000 kJ/mol nm^2^). All other degrees of freedom were unrestrained. This procedure allowed considerable conformational sampling while maintaining the N-SH2 domain in a particular conformational substate. The restraining potential was introduced gradually, while the temperature was increased from 50 to 300 K in 5 ns and then maintained constant for another 5 ns.

N-SH2 was pulled away from PTP using 600 ns pulling simulations at constant pull velocity (51, 52). Here, two sets of simulations were performed using two distinct reaction coordinates (RCs), respectively. First, RC_1_ was defined as the center-of-mass distance between the blocking loop (residues 60-62) and the catalytic PTP loop (residues 460-462). RC_1_ was pulled from 0.73nm to 2.15nm. Second, RC_2_ was defined as the distance between the C_α_ atoms of Asp^61^ and Ala^461^. RC_2_ was pulled from 0.55nm to 1.97nm. Only for free energy calculations along RC1, hydrogen atoms were described as virtual sites. To obtain statistically independent pathways for SHP2 opening, three pulling simulations were carried out for each reaction coordinate, each restrained either in the α-or in the β-state (12 simulations in total).

To further improve the conformation sampling during these pulling simulations and to obtain more independent starting frames for umbrella sampling (see below), we coupled the pulling simulations with simulated tempering (53). In line with previous studies (53, 54), simulated tempering simulations were carried out at constant volume. For simulated tempering, temperature steps of 5 K between 300 and 400 K were applied. Temperature changes were controlled according to the Metropolis algorithm (55) to obtain canonical ensembles at all temperatures. The initial weights were chosen following Park and Pande (54). Temperature transitions were attempted every 1 ps, and the weights were updated throughout the whole simulation according to the Wang-Landau adaptive weighting scheme (56). The pull force was set to 10000 kJ/mol nm^2^, whereas the reference position was changed with a velocity of 2.5×10^−6^ nm/ps.

### Free energy profiles of the SHP2 opening

The free energy profiles for the opening of SHP2 were obtained using umbrella sampling and the weighted histogram analysis method (WHAM) (57, 58). Four free energy profiles were computed: along RC_1_ or along RC_2_, and with N-SH2 either restrained in the α-or in the β-state. For each profile, 72 windows were used. Initial configurations for umbrella sampling were taken from the simulated tempering pulling simulations by means of a cluster analysis, thereby ensuring that the umbrella simulations were triggered from the most representative conformations. Accordingly, the configurations were collected from the sub-ensemble at 300K, divided in 72 groups on the base of the umbrella window interval they belong, and clustered using the GROMOS clustering method (59). The clustering cutoff was chosen on the basis of the root mean square deviation distribution in each group, picking the value corresponding to the first relative maximum in abscissa. The umbrella force constant was set to 4000 kJ mol^−1^nm^−2^, and sampling of 100 ns was performed for each window. Umbrella sampling simulations were carried out at constant pressure using the Parrinello-Rahman barostat (60). Statistical errors of the PMFs were estimated using the Bayesian bootstrap of complete histograms (58), thereby considering only complete histograms as independent.

### Simulations of the Thr^42^ mutants of the N-SH2 domain and opening of the β-strand

MD simulations were carried out on the N-SH2 domain in its apo form, considering five possible mutants that have been studied experimentally (6), arising from a single-base change at codon 42. These mutations involved the substitution of the Thr^42^ residue with Ala, Ser, Arg, Lys, or Ile. Starting from the wild-type structure, mutations were introduced with the Swiss-PdbViewer software package 4.1 (61) and the rotamers were chosen based on *(i)* favorable contacts with the protein and *(ii)* potential energies. The AMBER99SB force field was used. The equilibration procedure was the same as for wild-type N-SH2 domain. Production runs were performed for 500 ns. The structures were further equilibrated with a closed central β-sheet by first applying a harmonic restraint and then a holonomic constraint on the hydrogen bond between Gly^39^ and Asn^58^, forming a β-strand, while two simulated tempering simulations of 10 ns each were performed. Then, pulling simulations were carried out for gradually opening the β-strand over 250 ns. As RC, we here used the distance between the carbonyl C atom of Gly^39^ and the backbone N atom of Asn^58^. The pull force constant was set to 1000 kJ mol^−1^nm^−2^, whereas the reference Gly^39^ C–Asn^58^ N distance between the β-strands was changed with a velocity of 5×10^−6^ nm/ps. Again, simulated tempering (53) was used to enhance sampling, using a temperature range from 300 to 380 K in steps of 10 K. The free energy profiles were obtained using umbrella sampling and WHAM (57, 58). For each profile, 54 windows were used, the umbrella force constant was set to 1000 kJ mol^−1^nm^−2^, and a sampling of 500 ns was performed for each window. Statistical errors of the PMFs were estimated with the Bayesian bootstrap of complete histograms, yielding a maximum error of less than 2 kJ/mol (58).

### Effects of Thr^42^ substitutions on the propensity of N-SH2 for the α and β conformations

To test whether substitutions of Thr^42^ modulate the propensity of N-SH2 for the α- and β-states, we computed the free energies for mutating Thr^42^ with N-SH2 restrained to the α-or to β-state. These free-energy calculations were performed for the N-SH2 domain in its apo form or bound to the phosphopeptide SLNpYIDLDLVK. The restraint to α or β was again implemented with a restraint on a principal component (force constant 1000 kJ/mol nm^2^) (62). The free-energy calculations were carried out along an alchemical reaction coordinate λ. Thr^42^ in wild type N-SH2 (λ=0 state) was alchemically mutated to Ala, Ser, Arg, Lys and Ile, respectively (λ=1 state), The AMBER99SB force field was used, and the topologies for the alchemical transformation were generated using PMX (63). The free energy difference was computed using Crooks Fluctuation Theorem (31, 32, 63). Following the protocol in Ref. (63), each system was initially equilibrated, and slowly transformed from state λ=0 to state λ=1 within 10 ns. Next, equilibrium simulations for every mutation were performed for 10 ns for the states λ=0 and λ=1. From every equilibrium trajectory, 200 fast nonequilibrium runs of 100 ps each were spawned morphing the system from λ=0 to λ=1, and other 200 from λ=1 to λ=0. A soft-core potential was used for Lennard-Jones and Coulomb interactions (64). The statistical error was estimated by bootstrapping (sampling with replacement) from the fast non-equilibrium runs.

### Effects of peptide mutations on the affinity to the α-and β-states of N-SH2

To test if certain peptide mutations select for the α-or for the β-state of N-SH2, simulations were carried out on the N-SH2 domain in solution bound to SLNpYIDLDLVK or SPGEpYVNIEFGS, while restraining N-SH2 in the α-or β-state with a harmonic restraint on a principal component (k = 1000 kJ/mol nm^2^). Every substitution, with the exception of proline, was applied on these two phosphopeptides at the sequence position +5 and +6 relative to phosphotyrosine. Free-energy calculations were performed using Crooks Theorem and the procedure reported above, starting a set of five uncorrelated initial configurations obtained from independent equilibration simulations.

## ACKNOWLEDGEMENTS

This study has received funding from the European Union’s Horizon 2020 research and innovation programme under the Marie Skłodowska-Curie grant agreement MARS n. 705829. Early stages of this project were supported by Fondazione Umberto Veronesi (M.A.) and by the Deutsche Forschungsgemeinschaft (J.S.H., grant no 1971-4/1).

## REFERENCES

1 Blume-Jensen, P., & Hunter, T. (2001). Oncogenic kinase signalling. Nature, 411(6835), 355–365. doi:10.1038/35077225

2 Grossmann, K. S., Rosário, M., Birchmeier, C., & Birchmeier, W. (2010). Chapter 2 - The Tyrosine Phosphatase Shp2 in Development and Cancer. In G. F. Vande Woude & G. Klein (Eds.), Advances in Cancer Research (Vol. 106, pp. 53–89): Academic Press.

3 Tajan, M., de Rocca Serra, A., Valet, P., Edouard, T., & Yart, A. (2015). SHP2 sails from physiology to pathology. European Journal of Medical Genetics, 58(10), 509–525. doi:10.1016/j.ejmg.2015.08.005

4 Tartaglia, M., Mehler, E. L., Goldberg, R., Zampino, G., Brunner, H. G., Kremer, H., … Gelb, B. D. (2001). Mutations in PTPN11, encoding the protein tyrosine phosphatase SHP-2, cause Noonan syndrome. Nat Genet, 29(4), 465–468. doi:10.1038/ng772

5 Tartaglia, M., Martinelli, S., Stella, L., Bocchinfuso, G., Flex, E., Cordeddu, V., … Gelb, B. D. (2006). Diversity and functional consequences of germline and somatic PTPN11 mutations in human disease. Am J Hum Genet, 78(2), 279–290. doi:10.1086/499925

6 Martinelli, S., Torreri, P., Tinti, M., Stella, L., Bocchinfuso, G., Flex, E., … Tartaglia, M. (2008). Diverse driving forces underlie the invariant occurrence of the T42A, E139D, I282V and T468M SHP2 amino acid substitutions causing Noonan and LEOPARD syndromes. Hum Mol Genet, 17(13), 2018–2029. doi:10.1093/hmg/ddn099

7 Bocchinfuso, G., Stella, L., Martinelli, S., Flex, E., Carta, C., Pantaleoni, F., … Palleschi, A. (2007). Structural and functional effects of disease-causing amino acid substitutions affecting residues Ala72 and Glu76 of the protein tyrosine phosphatase SHP-2. Proteins, 66(4), 963–974. doi:10.1002/prot.21050

8 Digilio, M. C., Conti, E., Sarkozy, A., Mingarelli, R., Dottorini, T., Marino, B., … Dallapiccola, B. (2002). Grouping of multiple-lentigines/LEOPARD and Noonan syndromes on the PTPN11 gene. Am J Hum Genet, 71(2), 389–394. doi:10.1086/341528

9 Tartaglia, M., Martinelli, S., Iavarone, I., Cazzaniga, G., Spinelli, M., Giarin, E., … Biondi, A. (2005). Somatic PTPN11 mutations in childhood acute myeloid leukaemia. Br J Haematol, 129(3), 333–339. doi:10.1111/j.1365-2141.2005.05457.x

10 Tartaglia, M., Martinelli, S., Cazzaniga, G., Cordeddu, V., Iavarone, I., Spinelli, M., … Biondi, A. (2004). Genetic evidence for lineage-related and differentiation stage-related contribution of somatic PTPN11 mutations to leukemogenesis in childhood acute leukemia. Blood, 104(2), 307–313. doi:10.1182/blood-2003-11-3876

11 Loh, M. L., Martinelli, S., Cordeddu, V., Reynolds, M. G., Vattikuti, S., Lee, C. M., … Tartaglia, M. (2005). Acquired PTPN11 mutations occur rarely in adult patients with myelodysplastic syndromes and chronic myelomonocytic leukemia. Leuk Res, 29(4), 459–462. doi:10.1016/j.leukres.2004.10.001

12 Goemans, B. F., Zwaan, C. M., Martinelli, S., Harrell, P., de Lange, D., Carta, C., … Kaspers, G. J. (2005). Differences in the prevalence of PTPN11 mutations in FAB M5 paediatric acute myeloid leukaemia. Br J Haematol, 130(5), 801–803. doi:10.1111/j.1365-2141.2005.05685.x

13 Chen, Y. N., LaMarche, M. J., Chan, H. M., Fekkes, P., Garcia-Fortanet, Fortanet., Acker, M. G., … Fortin, P. D. (2016). Allosteric inhibition of SHP2 phosphatase inhibits cancers driven by receptor tyrosine kinases. Nature, 535(7610), 148–152. doi:10.1038/nature18621

14 Prahallad, A., Heynen, G. J., Germano, G., Willems, S. M., Evers, B., Vecchione, L., … Bernards, R. (2015). PTPN11 Is a Central Node in Intrinsic and Acquired Resistance to Targeted Cancer Drugs. Cell Rep, 12(12), 1978–1985. doi:10.1016/j.celrep.2015.08.037

15 Torres-Ayuso, Ayuso., & Brognard, J. (2018). Shipping Out MEK Inhibitor Resistance with SHP2 Inhibitors. Cancer Discov, 8(10), 1210–1212. doi:10.1158/2159-8290.CD-18-0915

16 Ahmed, T. A., Adamopoulos, C., Karoulia, Z., Wu, X., Sachidanandam, R., Aaronson, S. A., & Poulikakos, P. I. (2019). SHP2 Drives Adaptive Resistance to ERK Signaling Inhibition in Molecularly Defined Subsets of ERK-Dependent Tumors. Cell Reports, 26(1), 65–78.e65. doi:10.1016/j.celrep.2018.12.013

17 Hayashi, T., Senda, M., Suzuki, N., Nishikawa, H., Ben, C., Tang, C., … Hatakeyama, M. (2017). Differential Mechanisms for SHP2 Binding and Activation Are Exploited by Geographically Distinct Helicobacter pylori CagA Oncoproteins. Cell Rep, 20(12), 2876–2890. doi:10.1016/j.celrep.2017.08.080

18 Hui, E., Cheung, J., Zhu, J., Su, X., Taylor, M. J., Wallweber, H. A., … Vale, R. D. (2017). T cell costimulatory receptor CD28 is a primary target for PD-1-mediated inhibition. Science, 355(6332), 1428–1433. doi:10.1126/science.aaf1292

19 Butterworth, S., Overduin, M., & Barr, A. J. (2014). Targeting protein tyrosine phosphatase SHP2 for therapeutic intervention. Future Medicinal Chemistry, 6(12), 1423–1437. doi:10.4155/fmc.14.88

20 Ran, H., Tsutsumi, R., Araki, T., & Neel, B. G. (2016). Sticking It to Cancer with Molecular Glue for SHP2. Cancer Cell, 30(2), 194–196. doi:10.1016/j.ccell.2016.07.010

21 Frankson, R., Yu, Z.-H., Bai, Y., Li, Q., Zhang, R.-Y., & Zhang, Z.-Y. (2017). Therapeutic Targeting of Oncogenic Tyrosine Phosphatases. Cancer Research, 77(21), 5701. doi:10.1158/0008-5472.CAN-17-1510

22 Liu, B. A., & Machida, K. (2017). Introduction: History of SH2 Domains and Their Applications. Methods Mol Biol, 1555, 3–35. doi:10.1007/978-1-4939-6762-9_1

23 Hof, P., Pluskey, S., Dhe-Paganon, S., Eck, M. J., & Shoelson, S. E. (1998). Crystal structure of the tyrosine phosphatase SHP-2. Cell, 92(4), 441–450. doi:S0092-8674(00)80938-1

24 Lee, C. H., Kominos, D., Jacques, S., Margolis, B., Schlessinger, J., Shoelson, S. E., & Kuriyan, J. (1994). Crystal structures of peptide complexes of the amino-terminal SH2 domain of the Syp tyrosine phosphatase. Structure, 2(5), 423–438.

25 LaRochelle, J. R., Fodor, M., Vemulapalli, V., Mohseni, M., Wang, P., Stams, T., … Blacklow, S. C. (2018). Structural reorganization of SHP2 by oncogenic mutations and implications for oncoprotein resistance to allosteric inhibition. Nat Commun, 9(1), 4508. doi:10.1038/s41467-018-06823-9

26 Yu, Z. H., Xu, J., Walls, C. D., Chen, L., Zhang, S., Zhang, R., … Zhang, Z. Y. (2013). Structural and mechanistic insights into LEOPARD syndrome-associated SHP2 mutations. J Biol Chem, 288(15), 10472–10482. doi:10.1074/jbc.M113.450023

27 Barford, D., & Neel, B. G. (1998). Revealing mechanisms for SH2 domain mediated regulation of the protein tyrosine phosphatase SHP-2. Structure, 6(3), 249–254. doi:10.1016/S0969-2126(98)00027-6

28 Keilhack, H., David, F. S., McGregor, M., Cantley, L. C., & Neel, B. G. (2005). Diverse biochemical properties of Shp2 mutants. Implications for disease phenotypes. J Biol Chem, 280(35), 30984–30993. doi:10.1074/jbc.M504699200

29 Darian, E., Guvench, O., Yu, B., Qu, C. K., & MacKerell, A. D. Jr., (2011). Structural mechanism associated with domain opening in gain-of-function mutations in SHP2 phosphatase. Proteins, 79(5), 1573–1588. doi:10.1002/prot.22984

30 Anselmi, M., Calligari, P., Hub, J. S., Tartaglia, M., Bocchinfuso, G., & Stella, L. (2020). Structural Determinants of Phosphopeptide Binding to the N-Terminal Src Homology 2 Domain of the SHP2 Phosphatase. Journal of Chemical Information and Modeling. doi:10.1021/acs.jcim.0c00307

31 Crooks, G. E. (1998). Nonequilibrium Measurements of Free Energy Differences for Microscopically Reversible Markovian Systems. J. Stat. Phys., 90, 1481.

32 Crooks, G. E. (1999). Entropy production fluctuation theorem and the nonequilibrium work relation for free energy differences. Phys. Rev. E, 60, 2721.

33 Garcia Fortanet, J., Chen, C. H.-T., Chen, Y.-N. P., Chen, Z., Deng, Z., Firestone, B., … LaMarche, M. J. (2016). Allosteric Inhibition of SHP2: Identification of a Potent, Selective, and Orally Efficacious Phosphatase Inhibitor. Journal of Medicinal Chemistry, 59(17), 7773–7782. doi:10.1021/acs.jmedchem.6b00680

34 Gunasekaran, K., Ma, B., & Nussinov, R. (2004). Is allostery an intrinsic property of all dynamic proteins? Proteins, 57(3), 433–443. doi:10.1002/prot.20232

35 Chen, Y.-N. P., LaMarche, M. J., Chan, H. M., Fekkes, P., Garcia-Fortanet, J., Acker, M. G., … Fortin, P. D. (2016). Allosteric inhibition of SHP2 phosphatase inhibits cancers driven by receptor tyrosine kinases. Nature, 535(7610), 148–152. doi:10.1038/nature18621

36 Fedele, C., Ran, H., Diskin, B., Wei, W., Jen, J., Geer, M. J., … Tang, K. H. (2018). SHP2 Inhibition Prevents Adaptive Resistance to MEK Inhibitors in Multiple Cancer Models. Cancer Discov, 8(10), 1237–1249. doi:10.1158/2159-8290.CD-18-0444

37 Sun, X., Ren, Y., Gunawan, S., Teng, P., Chen, Z., Lawrence, H. R., … Wu, J. (2018). Selective inhibition of leukemia-associated SHP2E69K mutant by the allosteric SHP2 inhibitor SHP099. Leukemia, 32(5), 1246–1249. doi:10.1038/s41375-018-0020-5

38 Wong, G. S., Zhou, J., Liu, J. B., Wu, Z., Xu, X., Li, T., … Bass, A. J. (2018). Targeting wild-type KRAS-amplified gastroesophageal cancer through combined MEK and SHP2 inhibition. Nature Medicine, 24(7), 968–977. doi:10.1038/s41591-018-0022-x

39 Chemical Computing Group Inc. (2014). Molecular Operating Environment (MOE) software.

40 Jorgensen, W. L., Chandrasekhar, J., Madura, J. D., Impey, R. W., & Klein, M. L. (1983). Comparison of simple potential functions for simulating liquid water ‥ J. Chem. Phys., 79, 926–935.

41 Van Der Spoel, D., Lindahl, E., Hess, B., Groenhof, G., Mark, A. E., & Berendsen, H. J. (2005). GROMACS: fast, flexible, and free. J Comput Chem, 26(16), 1701–1718. doi:10.1002/jcc.20291

42 Hornak, V., Abel, R., Okur, A., Strockbine, B., Roitberg, A., & Simmerling, C. (2006). Comparison of multiple Amber force fields and development of improved protein backbone parameters. Proteins, 65(3), 712–725. doi:10.1002/prot.21123

43 Homeyer, N., Horn, A. H., Lanig, H., & Sticht, H. (2006). AMBER force-field parameters for phosphorylated amino acids in different protonation states: phosphoserine, phosphothreonine, phosphotyrosine, and phosphohistidine. J Mol Model, 12(3), 281–289. doi:10.1007/s00894-005-0028-4

44 Darden, T., York, D., & Pedersen, L. (1993). Particle Mesh Ewald: an Nlog(N) method for Ewald sum in large systems. J. Chem. Phys., 98, 10089–10092.

45 Miyamoto, S., & Kollman, P. A. (1992). SETTLE: An analytical version of the SHAKE and RATTLE algorithms for rigid water models . . J. Comp. Chem., 13, 952–962.

46 Hess, B., Bekker, H., Berendsen, H. J. C., & Fraaije, J. G. E. M. (1997). LINCS: a linear constraint solver for molecular simulations. Journal of Computational Chemistry, 18(12), 1463–1472.

47 Berendsen, H. J. C., Postma, J. P. M., Van Gunsteren, W. F., Di Nola, A., & Haak, J. R. (1984). Molecular dynamics with coupling to an external bath. Journal of Chemical Physics, 81(8), 3684–3690.

48 Bussi, G., Donadio, D., & Parrinello, M. (2007). Canonical sampling through velocity rescaling. J. Chem. Phys., 126, 14101.

49 Best, R. B., Zheng, W., & Mittal, J. (2014). Balanced Protein–Water Interactions Improve Properties of Disordered Proteins and Non-Specific Protein Association. Journal of Chemical Theory and Computation, 10(11), 5113–5124. doi:10.1021/ct500569b

50 Huang, J., Rauscher, S., Nawrocki, G., Ran, T., Feig, M., de Groot, B. L., … MacKerell, A. D., Jr. (2017). CHARMM36m: an improved force field for folded and intrinsically disordered proteins. Nat Methods, 14(1), 71–73. doi:10.1038/nmeth.4067

51 Izrailev, S., Stepaniants, S., Isralewitz, B., Kosztin, D., Lu, H., Molnar, F., … Schulten, K. (1998). Steered molecular dynamics Computational Molecular Dynamics: Challenges, Methods, Ideas (pp. 39–65): Springer-Verlag.

52 Grubmuller, H., Heymann, B., & Tavan, P. (1996). Ligand binding: molecular mechanics calculation of the streptavidin-biotin rupture force. Science, 271(5251), 997–999. doi:10.1126/science.271.5251.997

53 Marinari, E., & Parisi, G. (1992). Simulated tempering: a new Monte Carlo scheme. Europhys. Lett., 19, 451–458.

54 Park, S., & Pande, V. S. (2007). Choosing weights for simulated tempering. Phys Rev E Stat Nonlin Soft Matter Phys, 76(1 Pt 2), 016703. doi:10.1103/PhysRevE.76.016703

55 Metropolis, N., Rosenbluth, A. W., Rosenbluth, M. N., & Teller, A. H. (1953). Equation of state calculations by fast computing machines J. Chem. Phys., 21, 1087.

56 Wang, F., & Landau, D. P. (2001). Efficient, multiple-range random walk algorithm to calculate the density of states. Phys Rev Lett, 86(10), 2050–2053. doi:10.1103/PhysRevLett.86.2050

57 Kumar, S., Bouzida, D., Swendsen, S. H., Kollman, P. A., & Rosenberg, J. M. (1992). The Weighted Histogram Analysis Method for free-energy calculations on biomolecules. I. the method. Journal of Computational Chemistry, 13, 1011–1021.

58 Hub, J. S., de Groot, B. L., & van der Spoel, D. (2010). g_wham—A Free Weighted Histogram Analysis Implementation Including Robust Error and Autocorrelation Estimates. Journal of Chemical Theory and Computation, 6(12), 3713–3720. doi:10.1021/ct100494z

59 Daura, X., Gademann, K., Jaun, B., Seebach, D., van Gunsteren, W. F., & Mark, A. E. (1999). Peptide Folding: When Simulation Meets Experiment. Angewandte Chemie International Edition, 38(1 ‐ 2), 236–240. doi:doi:10.1002/(SICI)1521-3773(19990115)38:1/2<236::AID-ANIE236>3.0.CO;2-M

60 Parrinello, M., & Rahman, A. (1981). Polymorphic transitions in single crystals: A new molecular dynamics method. J. Appl. Phys., 52, 7182–7190.

61 Guex, N., & Peitsch, M. C. (1997). SWISS-MODEL and the Swiss-PdbViewer: an environment for comparative protein modeling. Electrophoresis, 18(15), 2714–2723. doi:10.1002/elps.1150181505

62 Lange, O. E., Schafter, L. V., & Grubmuller, H. (2006). Flooding in GROMACS: accelerated barrier crossing in molecular dynamics. Journal of Computational Chemistry, 27, 1693–1702.

63 Gapsys, V., Michielssens, S., Seeliger, D., & de Groot, B. L. (2015). pmx: Automated protein structure and topology generation for alchemical perturbations. J Comput Chem, 36(5), 348–354. doi:10.1002/jcc.23804

64 Beutler, T. C., Mark, A. E., van Schaik, R. C., Greber, P. R., & van Gunsteren, W. F. (1994). Avoiding singularities and numerical instabilities in free energy calculations based on molecularsimulations. Chem. Phys. Lett., 222, 529–539.

