## Supplemental information for "An allosteric interaction controls the activation mechanism of SHP2 tyrosine phosphatase"

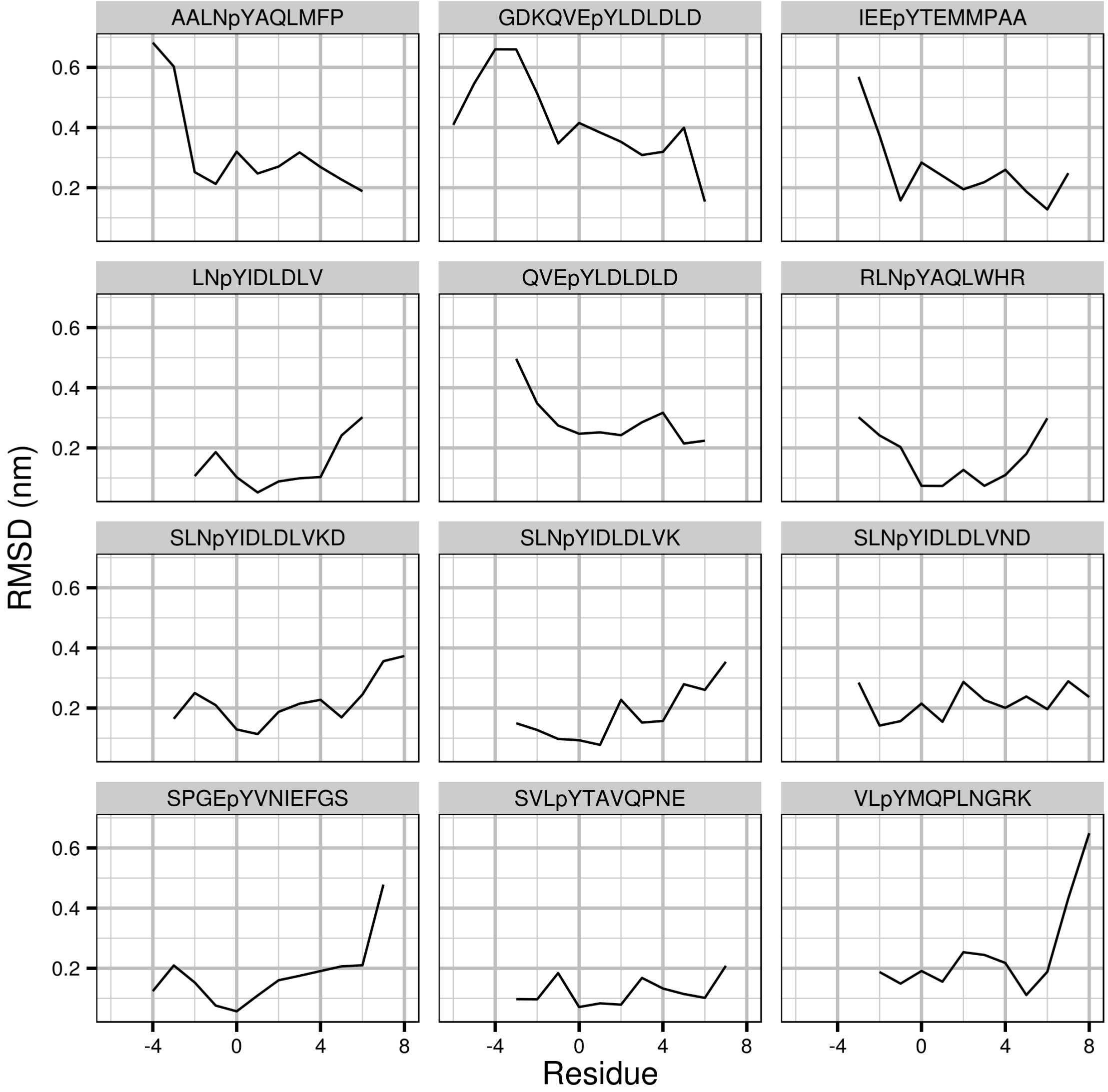


**Figure S1.** Root mean-square deviation (RMSD) of the most representative MD conformation defined as the central structure of the main structural cluster. The RMSD of the phosphopeptide $C_\alpha$ atoms is given relative the corresponding reference crystal structure, after superimposing only the backbone atoms of the N-SH2 domain.

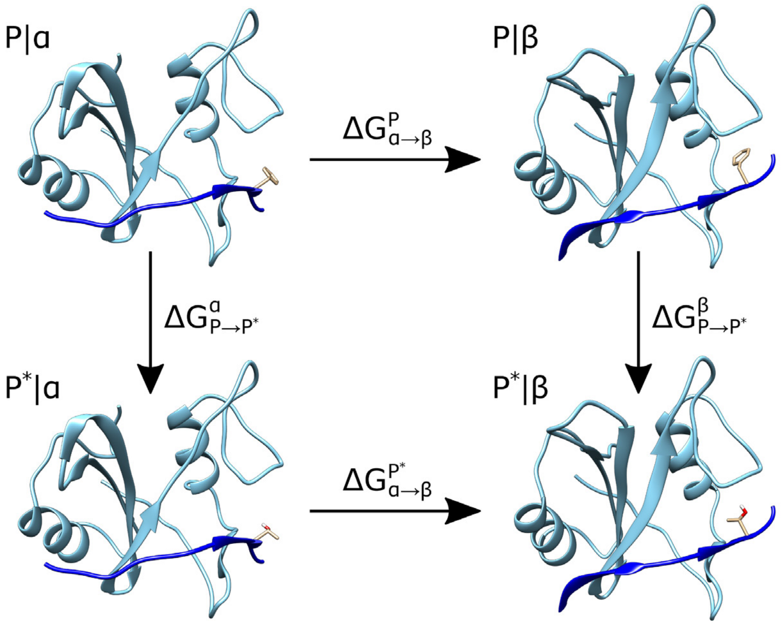


**Figure S2.** Thermodynamic cycle used for computing the change of preference of the ligand/N-SH2 complex for the α or the β state upon introducing a mutation in the ligand. Change of the stability of the ligand/N-SH2 complex upon introducing a mutation in the ligand, given by the respective free energy difference $\Delta G_{P\rightarrow P*}$, where $P$ and $P*$ denote the initial and the alchemically transformed amino acid, respectively. To quantify the effect of the $P \rightarrow P^*$ mutation on the N-SH2 preference for the α versus the β state, we computed $\Delta G_{P\rightarrow P*}$, twice for each mutation, namely with N-SH2 restrained either in the α or in the β state, denoted $\Delta G^{\alpha}_{P\rightarrow P*}$ and $\Delta G^{\beta}_{P\rightarrow P*}$ , respectively. Hence, the difference $\Delta\Delta G = \Delta G^{\beta}_{P\rightarrow P*} - \Delta G^{\alpha}_{P\rightarrow P*}$ indicates the change of preference of the ligand/N-SH2 complex for the α or the β state, where a positive $\Delta\Delta G$ indicates an augmented preference for the α state, and a negative $\Delta\Delta G$ an augmented preference for the β state. Critically, the $\Delta\Delta G$ values represent purely the change of the α-versus-β population for a bound ligand.

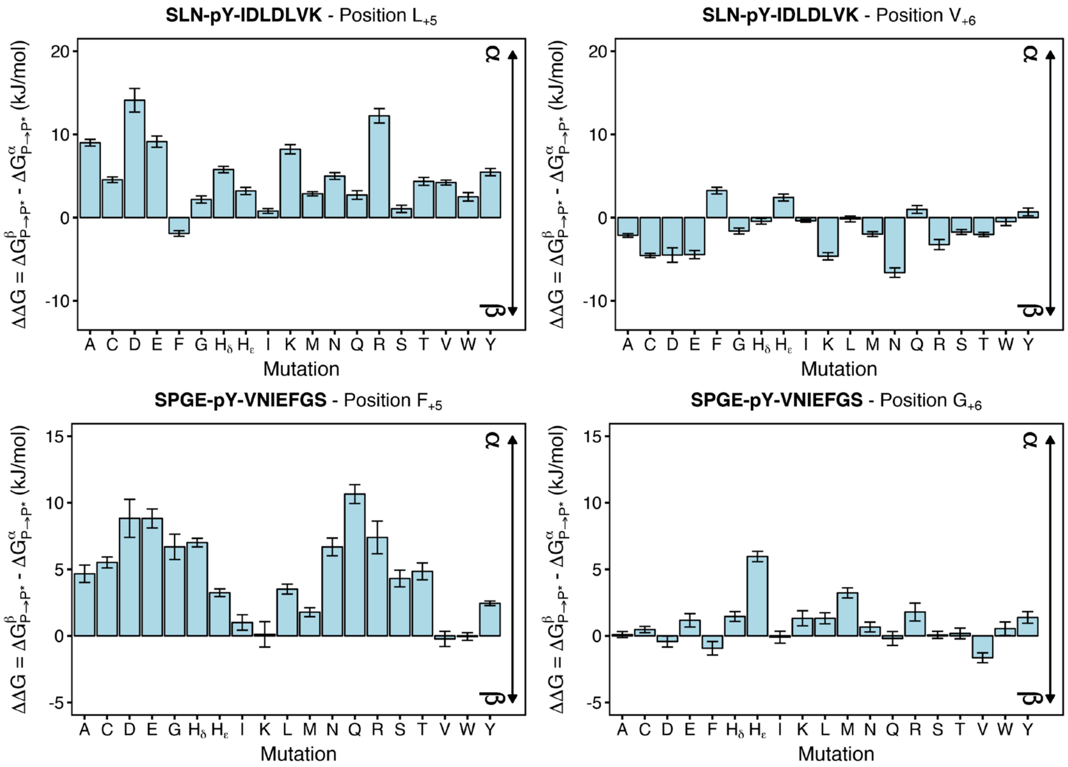


**Figure S3.** Change of preference of the ligand/N-SH2 complex for the α or the β state upon introducing a mutation in the ligand. Positive values of ΔΔG indicates an augmented preference for the α state respect to the reference sequence, and a negative Δ*G* an augmented preference for the β state. The calculations were performed on the position +5 and +6 of two ligands, SLNpYIDLDLVK and SPGEpYVNIEFGS, yielding the analogs SLNpYIDLDX$_{+5}$VK, SLNpYIDLDLX$_{+6}$K, SPGEpYVNIEX$_{+5}$GS and SPGEpYVNIEFX$_{+6}$S, where X considers all possible substitutions with the exception of proline.

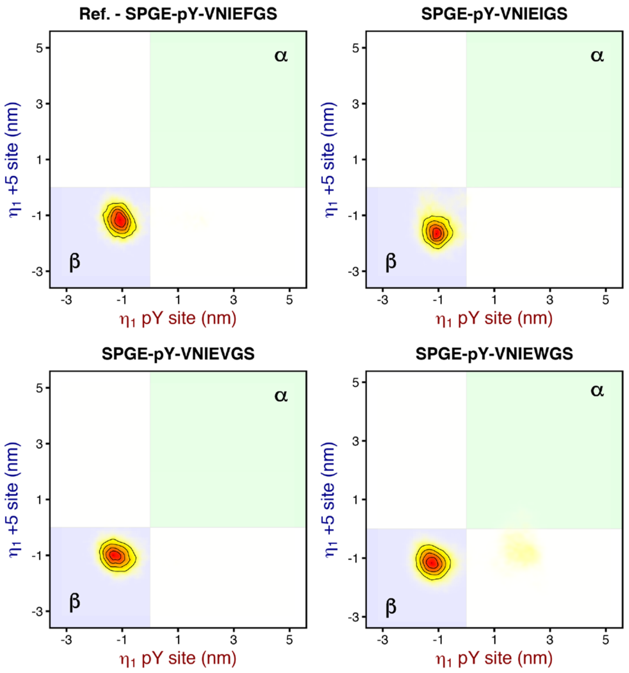


**Figure S4.** Projection distributions of the trajectory of the N-SH2 domain bound to the reference SPGEpYVNIEFGS peptide and its mutants at position +5, SPGEpYVNIE$X_{+5}$GS (X = I, V, W), on the principal component plane defined by the principal component eigenvector describing the motion of the pY loop and of the +5 site. The region of the plane corresponding to the α state (pY loop closed, +5 site closed) is shaded in green, whereas that of the β state (pY loop open, +5 site open) is shaded in blue. The substitution of the phenylalanine residue at position +5 in SPGEpYVNIEFGS respectively in isoleucine (I), valine (V), and tryptophan (W) retains the peptide binding mode to select the β state.

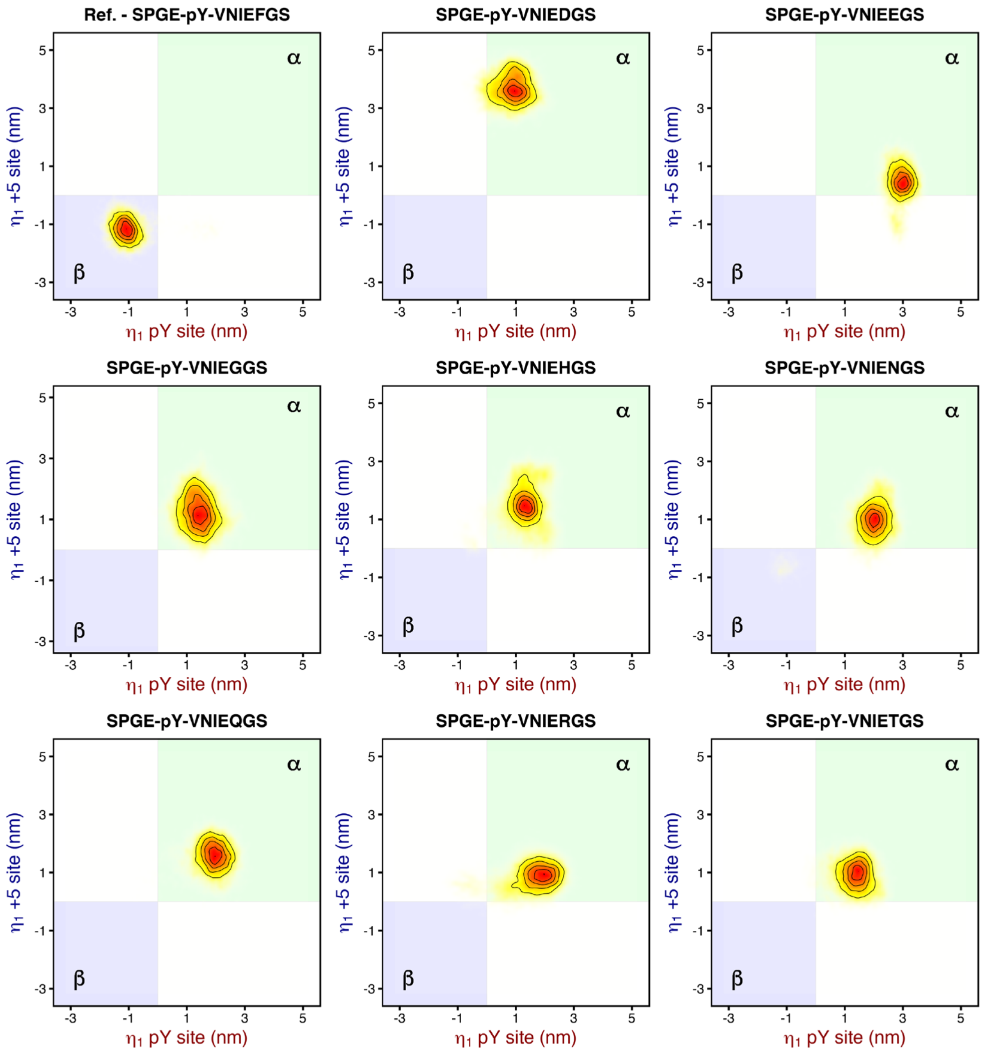


**Figure S5.** Projection distributions of the trajectory of the N-SH2 domain bound to the reference SPGEpYVNIEFGS peptide and its mutants at position +5, SPGEpYVNIEX$_{+5}$GS (X = D, E, G, H, N, Q, R, T), on the principal component plane defined by the principal component eigenvector describing the motion of the pY loop and of the +5 site. The region of the plane corresponding to the α state (pY loop closed, +5 site closed) is shaded in green, whereas that of the β state (pY loop open, +5 site open) is shaded in blue. The substitution of the phenylalanine residue at position +5 in SPGEpYVNIEFGS respectively in aspartate (D), glutamate (E), glycine (G), histidine (H$_\delta$), asparagine (N), glutamine (Q), arginine (R), and threonine (T) turns the peptide binding mode from selecting the β state to favoring the α state.

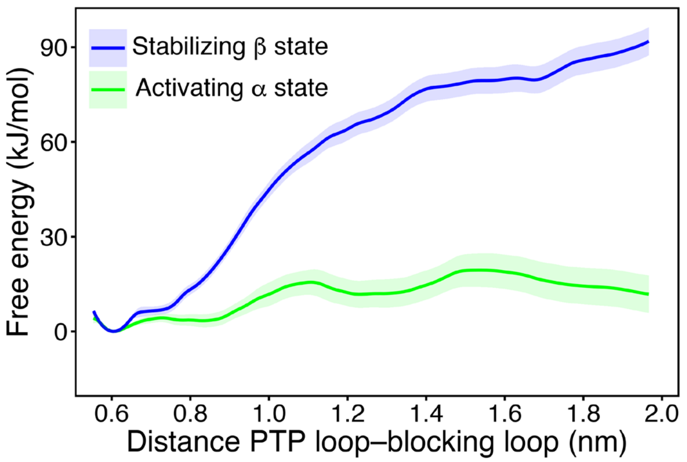


**Figure S6.** Free energy profiles for the opening of SHP2 using an alternative reaction coordinate ($RC_2$), with the N-SH2 domain restrained in the activating α state (green line) or in the stabilizing β state (blue line). $RC_2$ was defined as the distance between the $C_\alpha$ atoms of residues $Asp^{61}$ and $Ala^{461}$. The profiles are qualitatively similar to the profiles computed with $RC_1$.

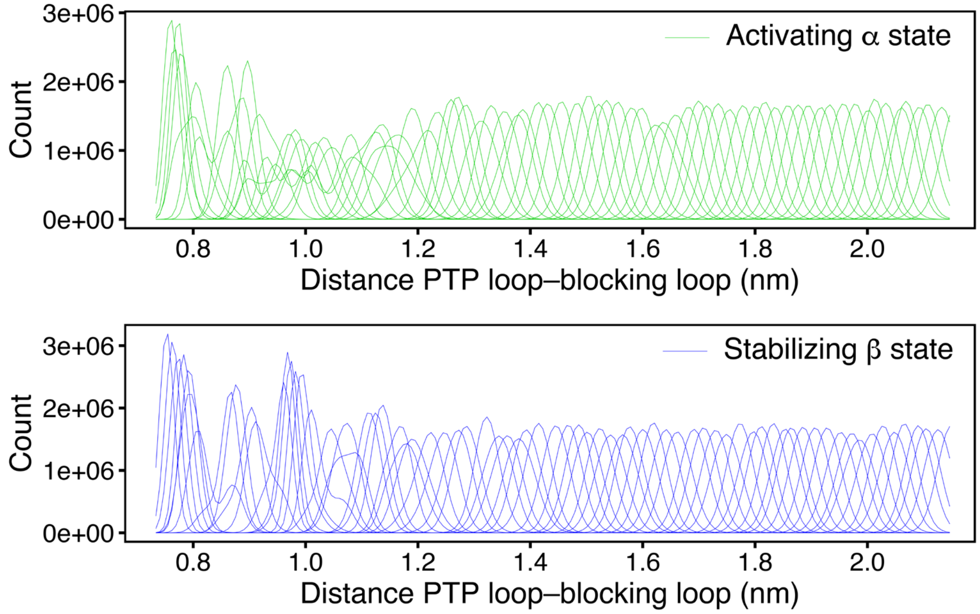


**Figure S7.** Umbrella histograms used for calculating of free energy profiles along the reaction coordinate $RC_1$, with the N-SH2 domain restrained in the activating α state (green lines) or in the stabilizing β state (blue lines). Evidently, the histograms well overlap, as required for reliable free energy profile calculations.

**Table S1. Degree of openness of the central beta strands βC and βD in various crystal structures, quantified by the distance between the backbone carbonyl oxygen of Gly$^{39}$ and the backbone nitrogen atom of Asn$^{58}$. The structures of isolated N-SH2 bound to peptides are shaded in green. The structures of wild type SHP2 and mutants are shaded in light and dark blue respectively. Other structures are shaded in yellow.**

| Pdb ID | Distance GLY$^{39}$O-ASN$^{58}$N (Å) | Description | Notes |
|---|---|---|---|
| 1AYA | 7.225 | N-SH2 only / bound to high affinity peptide | |
| 1AYB | 6.611 | N-SH2 only / bound to high affinity peptide | |
| 1AYC | 7.558 | N-SH2 only / bound to low affinity peptide | |
| 3TL0 | 7.679 | N-SH2 only / bound to low affinity peptide | |
| 4QSY | 7.356 | N-SH2 only / bound to low affinity peptide | |
| 2SHP | 3.722 | SHP2 | |
| 4DGP | 3.451 | SHP2 | |
| 4DGX | 3.499 | SHP2 Y279C mutant | |
| 4GWF | 4.020 | SHP2 Y279C mutant | BG loop incomplete |
| 4H1O | 4.749 | SHP2 D61G mutant | BG loop incomplete |
| 4H34 | 4.803 | SHP2 Q506P mutant | BG loop incomplete |
| 4NWF | 3.098 | SHP2 N308D mutant | BG loop incomplete |
| 4NWG | 4.877 | SHP2 E139D mutant | BG loop incomplete |
| 4OHD | 2.956 | SHP2 A461T mutant | |
| 4OHE | 4.322 | SHP2 G464A mutant | |
| 4OHH | 4.112 | SHP2 Q506P mutant | |
| 4OHI | 4.333 | SHP2 Q510E mutant | |
| 4OHL | 4.116 | SHP2 T468M mutant | |
| 5I6V | 3.042 | SHP2 F285S mutant | BG loop incomplete |
| 5IBM | 3.477 | SHP2 S502P mutant | BG loop incomplete |
| 5IBS | 3.755 | SHP2 E76Q mutant | BG loop incomplete |
| 1AYD | 5.244 | N-SH2 only / unbound | |
| 4JE4 | 5.207 | Crystal Structure of Monobody NSa1/SHP2 N-SH2 Domain Complex | |
| 5DF6 | 4.947 | tandem SH2 domains in complex with a TXNIP peptide | |
| 5EHP | 4.429 | SHP2 in Complex with Allosteric Inhibitor SHP836 | PO4 bound |
| 5EHR | 4.455 | SHP2 in Complex with Allosteric Inhibitor SHP099 | PO4 bound |

**Table S2. Modeling of initial conformations from PDB structures**

Initial structures of N-SH2/ligand complexes were generated from crystal structures with PDB codes listed in the table. To this end, PDB structures with similar ligand sequence were modified by means of Molecular Operative Environment (MOE), followed by a local energy minimization with side chain repacking and visual analysis. The substituted residues are highlighted in red.

| Simulated sequence | Original Sequence | PDB code |
|---|---|---|
| SLNpYIDLDLVK | GDKQVEpYLDLDLD | 4qsy |
| LNpYIDLDLV | GDKQVEpYLDLDLD | 4qsy |
| AALNpYAQLMFP | SVLpYTAVQPNE | 1aya |
| SPGEpYVNIEFGS | SPGEpYVNIEFGS | 1ayb |
| VLpYMQPLNGRK | SVLpYTAVQPNE | 1aya |
| IEEpYTEMMPAA | SVLpYTAVQPNE | 1aya |
| SVLpYTAVQPNE | SVLpYTAVQPNE | 1aya |
| RLNpYAQLWHR | RLNpYAQLWHR | 3tl0 |
| GDKQVEpYLDLDLD | GDKQVEpYLDLDLD | 4qsy |
| QVEpYLDLDLD | GDKQVEpYLDLDLD | 4qsy |
